# Cell-Free Characterization of Coherent Feed-Forward Loop-Based Synthetic Genetic Circuits

**DOI:** 10.1101/2021.01.11.426179

**Authors:** Pascal A. Pieters, Bryan L. Nathalia, Ardjan J. van der Linden, Peng Yin, Jongmin Kim, Wilhelm T.S. Huck, Tom F. A. de Greef

## Abstract

Regulatory pathways inside living cells employ feed-forward architectures to fulfill essential signal processing functions that aide in the interpretation of various types of inputs through noise-filtering, fold-change detection and adaptation. Although it has been demonstrated computationally that a coherent feed-forward loop (CFFL) can function as noise filter, a property essential to decoding complex temporal signals, this motif has not been extensively characterized experimentally or integrated into larger networks. Here we use post-transcriptional regulation to implement and characterize a synthetic CFFL in an *Escherichia coli* cell-free transcription-translation system and build larger composite feed-forward architectures. We employ microfluidic flow reactors to probe the response of the CFFL circuit using both persistent and short, noise-like inputs and analyze the influence of different circuit components on the steady-state and dynamics of the output. We demonstrate that our synthetic CFFL implementation can reliably repress background activity compared to a reference circuit, but displays low potential as a temporal filter, and validate these findings using a computational model. Our results offer practical insight into the putative noise-filtering behavior of CFFLs and show that this motif can be used to mitigate leakage and increase the fold-change of the output of synthetic genetic circuits.

## Introduction

A multitude of critical biological functions in cells, such as growth and differentiation, are regulated by dedicated genetic circuits.^1,2^ Synthetic equivalents of these circuits have been developed, including toggle switches,^3^ oscillators^3^ and logic gates,^4^ which has driven the development of more complex synthetic modules. For example, bistable switches have been utilized to create complex finite-state machines,^5^ genetic AND-logic gates have been employed to generate synthetic T cell-based therapies,^6,7^ and a feedback motif has been used to create a multi-layered cell structure through a synthetic differentiation circuit.^8^ Nevertheless, the topology of a genetic circuit does not necessarily dictate a single unique function, since circuits can have multiple functions depending on variations in parameters^9 –11^ and network motifs in general can produce diverse behavior,^12^ with only some constraints posed by their topologies.^13,14^ It is therefore crucial to construct and study genetic circuits to elucidate their range of functions and their behavior in larger networks.

When implementing synthetic networks of increasing sizes, undesired interactions with host organism machinery and excessive load on the host can impede the function of the synthetic circuit.^15^ The use of *in vitro* transcription and translation (TXTL) reactions eliminates the need for a host organism and provides a biomolecular breadboarding environment to rapidly construct synthetic genetic networks.^16–19^ TXTL reactions provide a flexible environment to dynamically vary inputs and circuit parameters without the need for extensive bacterial cloning and culture cycles.^17^ A toolbox of genetic elements has been created through extensive characterization of the TXTL reaction mixture alongside *E. coli* and phage-derived transcriptional regulators.^16,19^ This TXTL toolbox has been successfully utilized to rapidly construct and study gene cascades,^16^incoherent feed-forward loops,^16,20^ a negative feedback loop^21^ and oscillators.^22–25^ Although the toolbox extends the range of available genetic elements through inclusion of native *E. coli* transcription factors, which would severely interfere with an *E. coli* host cell when utilized *in vivo*, the number of available regulatory elements is still limited and does not scale up easily for the construction of topologically complex genetic networks. To resolve this, an additional regulatory layer can be introduced by utilizing post-transcriptional interactions such as riboswitches. In bacteria, small RNAs form a class of regulators that extend the complexity of genetic networks beyond transcriptional regulation.^26^ For synthetic circuits, toehold switch riboregulators, which can be forward-engineered, provide a wide dynamic range and are highly orthogonal, offer the potential to construct larger synthetic genetic networks.^27–29^

Here, we successfully construct a modular synthetic gene network based on translational regulation using toehold switch riboregulators in TXTL. We design and build a coherent feed-forward loop (CFFL) with RNA as top regulator, modeled after sRNA-based CFFLs found in nature (Figure 1a).^30–33^ The CFFL is a network motif highly abundant in both bacterial and mammalian regulatory networks^34^ and can display temporal filtering through sign-sensitive delay, where short inputs do not provoke a response while long-lived, persistent inputs are capable of generating a strong response.^35–37^ This property is essential for circuits experiencing a noisy input signal but can also serve to decode the temporal information that is encoded in various stimuli in cells.^38–42^ However, due to the dynamic nature of this function, it has remained difficult to systematically analyze temporal properties of CFFLs, with experimental characterization being limited to a narrow subset of its behavior.^37^ Next to temporal filtering, it has been postulated that CFFLs can suppress leaky expression in a network, resulting in a higher fold-change of the circuit output.^30^

**Figure 1:**
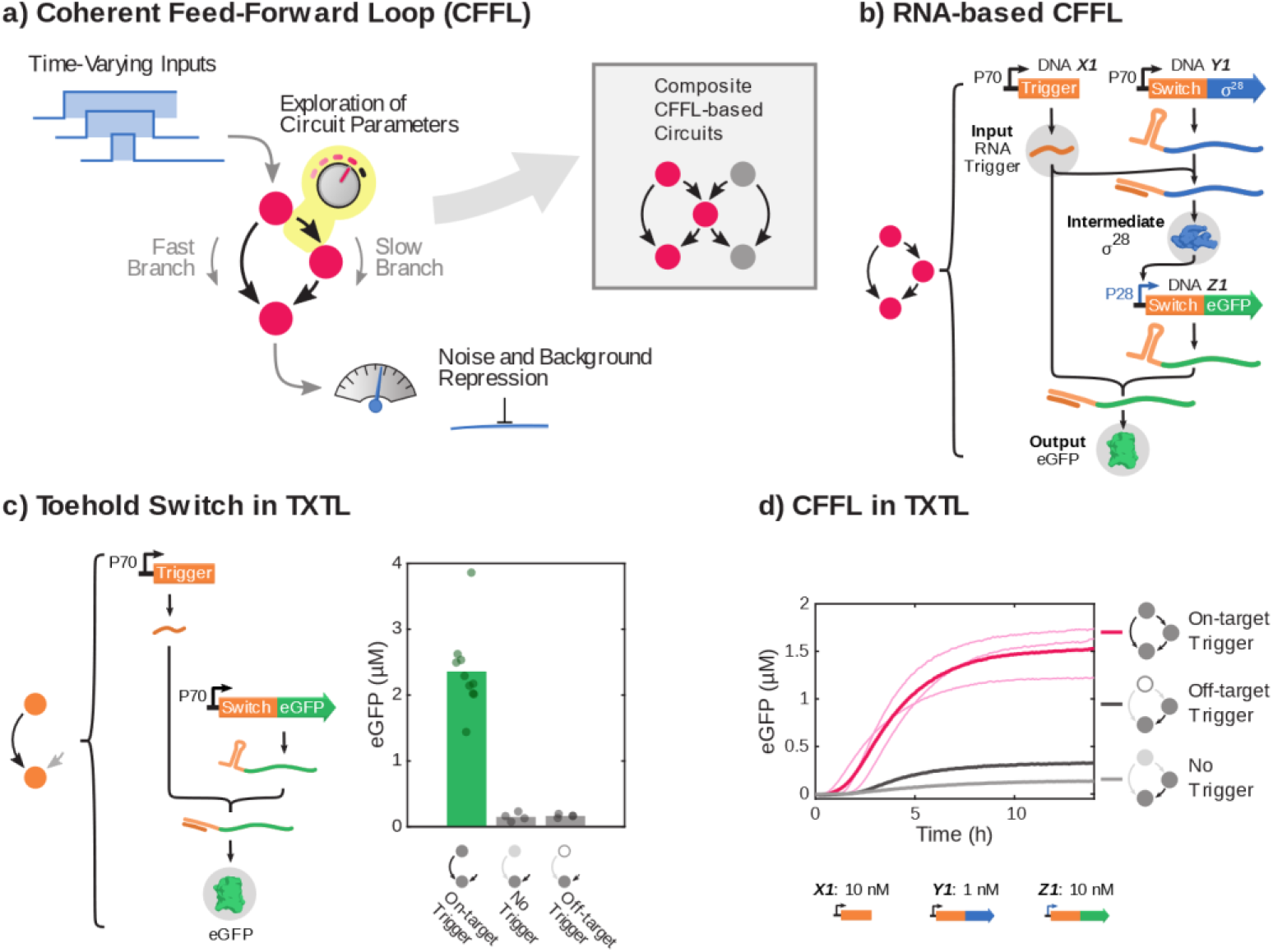
General concept of the construction and characterization of synthetic circuits based on CFFLs. a) Schematic drawings, where nodes are genes or RNA genes and arrows indicate interactions, of a CFFL and a composite network consisting of two CFFLs joined on their intermediate node (inset). b) Schematic representation of all DNA species and the DNA, RNA and protein level interactions that constitute the CFFL. The three main components are the toehold switch, marked by the 5’-adjacent RNA stem loop, and its corresponding RNA trigger (orange), the E. coli σ^28^-factor (blue) and fluorescent output protein (green). c) The toehold switch, which requires a matching RNA trigger to unfold the stem loop that obscure the RBS of a gene, was expressed in TXTL reactions. Endpoint eGFP concentrations (after 14h of incubation) for the toehold switch DNA construct (2 nM) in the presence of on-target trigger DNA (10 nM), off-target trigger DNA (10 nM) and without trigger. d) Time traces of eGFP production for the CFFL (where light pink traces are three distinct experiment and dark pink is their average). Expression without a trigger construct (light gray) and with an off-target trigger (dark gray) are plotted as negative controls. All DNA constructs and concentrations are summarized in Table S3.

We use the modularity of our design to implement several CFFL variants with orthogonal post-transcriptional toehold switch-based regulation^27^and, inspired by naturally occurring feed-forward architectures,^43^ combine the CFFL variants into a topologically more complex feed-forward circuit. We characterize both the background suppression and temporal filtering functions of the synthetic CFFL circuit using a microfluidic semi-continuous flow reactor to sustain prolonged TXTL reactions,^23–25,44^ complemented with *in silico* experiments. Our analysis reveals that the synthetic CFFL can effectively reduce background expression of components, increasing the fold-change of circuits for a wide range of circuit parameters. In agreement with recent computational studies that show low robustness of temporal filtering in CFFLs,^35^ the synthetic CFFL is not a potent noise filter, since the response of the circuit to time-varying inputs is similar to a reference cascade. We identify that more ultrasensitive response is required in the circuit component responsible for the delay in signal propagation, in order to develop a CFFL circuit that is able to serve as noise filter. Our results provide a foundation to construct modular synthetic gene networks based on translational regulation, demonstrated by the creation of complex CFFL-based circuits, and offer renewed insight into the signal processing functions of feed-forward loops.

## Results

The structurally simplest CFFL motif consists of three genes that interact to enable a signal to propagate from the input either directly or via an intermediate gene to the output gene (Figure 1a). In this circuit, the dissimilarity between the two pathways allows for propagation of the signal with different timescales. The delay generated by the presence of an intermediate gene largely determines the difference in timescales, whereas the mechanism by which the two pathways are integrated controls which aspect of the output is governed by the induced delay.^37^

### In vitro implementation of a synthetic CFFL

We designed a synthetic type 1 coherent feed-forward loop, representing the most commonly observed subtype of CFFLs,^36^ with AND-gate logic integrating the two branches of the circuit. In our design, the *E. coli* σ^28^-factor, which has been successfully used in various TXTL-based genetic circuits,^22,25^ is employed as intermediate species. The highly programmable toehold switch and trigger RNA-RNA post-transcriptional interactions are used to implement AND-type behavior (Figure 1b).^27^ Additionally, the RNA trigger is utilized to activate translation of the sigma-factor, creating a CFFL with an RNA species as a top regulator, mimicking naturally occurring RNA regulatory circuits.^30,32^

Genes for the synthetic CFFL were constructed using a Golden Gate assembly-based cloning method,^17^ enabling rapid prototyping of various promoter, toehold switch, and protein combinations (Figure S1). First, fragments of the CFFL were constructed and characterized in isolation. When a toehold switch and trigger pair were expressed in TXTL, we initially observed high background expression, but were able to largely eliminate this by optimizing RNA stability and removing in-frame start codons not regulated by the toehold switch (Figure S2). Batch expression from the toehold switch construct in the presence of the cognate RNA trigger is comparable to expression from an eGFP reference construct with a highly efficient ribosome binding site (RBS), whereas background expression in the absence of trigger is an order of magnitude lower (Figure 1c; Figure S3). As expected, expression increased monotonically for increasing concentrations of both switch and trigger DNA.

We assessed whether expression from a P70a promoter was impacted by competition between the *E. coli* sigma factor σ^28^ and the housekeeping σ^70^-factor for the RNA polymerase, but found no decrease in expression level from the constitutive promoter (Figure S4). The toehold switch was subsequently utilized as a translational regulator for σ^28^, resulting in a cascade that constitutes one branch of the CFFL. Upon expression in batch TXTL reactions, we observed significant increase in cascade activation only when on-target RNA trigger was produced (Figure S5). Next, this cascade was extended to contain a toehold switch in the 5’-UTR of the eGFP output construct, resulting in the synthetic CFFL motif. When expressing the CFFL constructs in TXTL, eGFP expression only increased drastically in the presence of the cognate RNA trigger (Figure 1d). In the absence of trigger, some leakage from the switch was observed and slight crosstalk with a randomly selected off-target trigger was detected. In conjunction with the CFFL, a reference motif was designed by removing the interaction between the input RNA trigger and the intermediate σ^28^-factor construct (Figure S6a). This circuit represents a simple signal transducing network with only a single path from input to output, to which the CFFL circuit characteristics can fairly be compared.^36^ Similar to the CFFL, the reference motif was confirmed to only activate when the DNA construct of the on-target RNA trigger was present in the TXTL reactions (Figure S6b). These circuits collectively constitute a flexible system to build and analyze CFFL-based networks.

### Composite CFFL circuits

Feed-Forward loops are often organized in constitutions of multiple interconnected loops of different architectural principles that can have distinct information processing functions.^43^ To demonstrate the feasibility of implementing these CFFL-based circuits with increased topological complexity in TXTL reactions, we constructed two new CFFL variations (CFFL 2 and 3) based on our initial design (CFFL 1) and merged them into a composite 5-node CFFL design (Figure 2a). First, we implemented an orthogonal CFFL circuit (CFFL 2) by replacing the toehold switch and trigger (switch/trigger A) of the initial design with a largely orthogonal switch and trigger pair (switch/trigger B), selected from two switch/trigger pairs evaluated for their dynamic range in TXTL. Secondly, we constructed a CFFL variant with eCFP as output protein while maintaining the switch/trigger A combination (CFFL 3). Both alternative implementations achieved comparable expression levels and retained a clear distinction between on and off states (Figure 2b).

**Figure 2:**
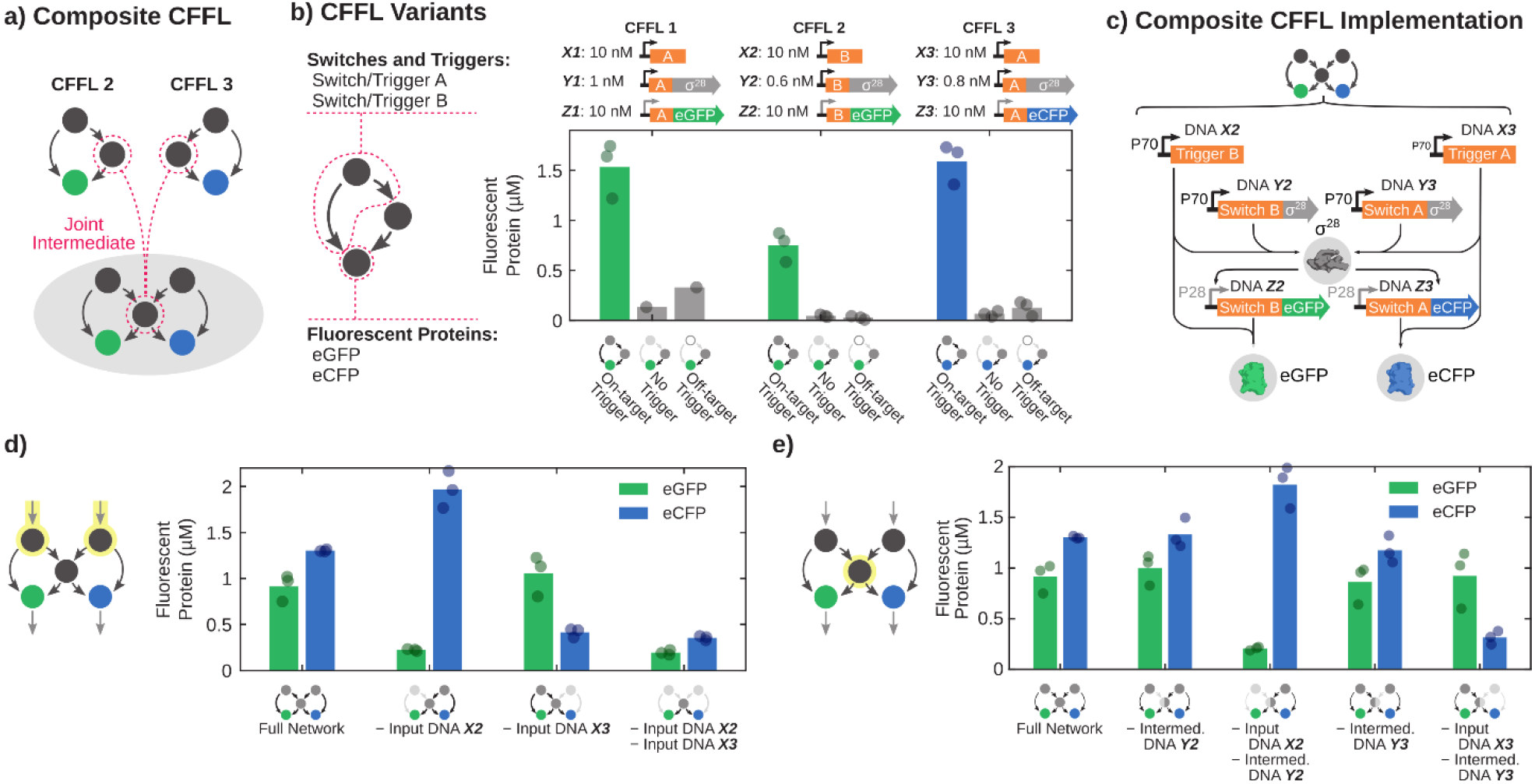
Construction of a composite CFFL using new synthetic CFFL variants. a) Schematic drawing of the formation of a composite CFFL that shares the intermediate σ^28^-factor protein (middle), using two new CFFL variants (CFFL 2 and 3). b) Endpoint fluorescent protein concentrations (after 14h incubation) of CFFL (CFFL 1), a variant with a distinct switch (CFFL 2) and a CFFL with eCFP as output protein (CFFL 3). c) Schematic representation of all DNA species and the DNA, RNA and protein level interactions that form the composite CFFL with a shared intermediate σ^28^-factor protein (middle), using CFFL 2 (left) and CFFL 3 (right). d) Endpoint eGFP and eCFP concentrations of the composite CFFL with both input DNA constructs present, with either of the inputs or without input. e) Endpoint concentrations for the composite CFFL when one of the σ^28^-producing DNA constructs is omitted. Concentrations of DNA species in d) and e) are as shown in b). In case an off-target trigger was used, its DNA concentration was 10 nM, equal to the DNA concentration for on-target triggers. All DNA constructs and concentrations are summarized in Table S3.

Using CFFL variant 2 and 3, we constructed a composite CFFL with two inputs and outputs and with the σ^28^ gene as a shared intermediate node (Figure 2c). Although not explored here, the design of the CFFL system allows for the modular substitution of the intermediate σ^28^ gene by other σ-factors^16^ to create the remaining composite CFFLs described by Gorochowski *et al.*^43^ with distinct intermediate genes. We evaluated the composite CFFL by expressing all DNA constructs of CFFL 2 and CFFL 3 (Figure 2d) in a single TXTL reaction whilst monitoring the eGFP and eCFP output fluorescence. The observed expression levels mostly equaled the individual CFFL circuits, ruling out large contributions of the synergistic production of σ^28^ and depletion of resources.^45^ When we omitted either one of the input triggers, expression of the corresponding fluorescent protein dropped significantly, while omission of both inputs yielded background levels of all outputs, indicating that the output RNA constructs are correctly activated by their cognate triggers. The σ^28^ protein that serves as intermediate for both sides of the composite circuit was produced in excess, since removal of either of the σ^28^-encoding DNA constructs did not lower the output levels of the circuit (Figure 2e). Nevertheless, the production of σ^28^ by one of the inputs can drive the production of RNA for the opposite output construct, as revealed by combinatorial evaluation of circuit components (Figure S7). In summary, we demonstrate that composite feed-forward organizations can be readily implemented using our synthetic CFFL design and the composite CFFL with a shared intermediate node displayed selective activation of each output by its cognate input while simultaneously being coupled to the opposite input.

### Characteristics of the synthetic CFFL and composite CFFL

We have demonstrated that the synthetic CFFL can propagate an input signal and can be used to implement topologically more complex feed-forward circuits. To assess if the circuit can display information processing functionalities associated with a CFFL network motif^30,36,37^ and whether these functionalities are retained in the composite CFFL network, we observed the CFFL, reference motif, and composite CFFL over a range of circuit parameters. A range of relative expression levels of the circuit components was sampled by varying the concentration of the DNA species. The response of the circuits was analyzed using four circuit characteristics (Figure 3a). Firstly, to quantify the repression of background expression by the CFFL-based circuits, *ON* and *OFF* state expression levels were determined. Additionally, the ratio between these two measures, the *ON/OFF* ratio, was computed as a measure of the relative change in output upon circuit activation. TXTL batch reactions are unsuitable for the full assessment of temporal filtering behavior of the synthetic CFFL due to the inability to apply time-varying inputs. Nevertheless, a characteristic time scale of the response of the circuit can be determined and used to estimate for which range of input pulse durations the behavior is expected to manifest. We therefore computed, *t_50_*, the time until half of the maximum output was reached as a measure of the characteristic circuit time-scale.

**Figure 3:**
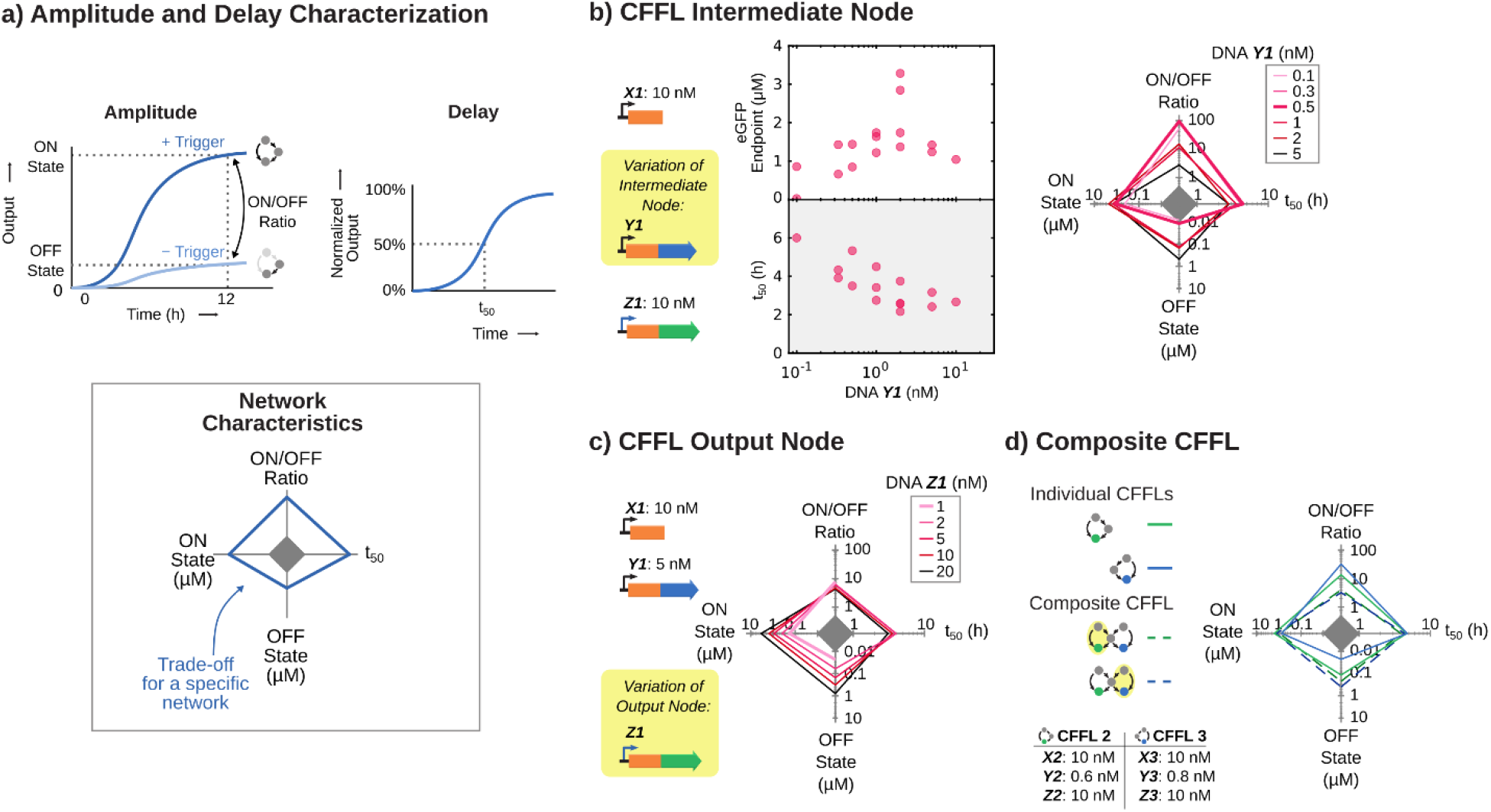
Endpoint and kinetic characterization of the CFFL implemented using TXTL batch reactions. a) Schematic representation of the four characteristics determined for each circuit and parameter combination, plotted on the axes of a spider plot. The endpoint concentration of the output protein with circuit input (ON state; left axis) or without (OFF state; bottom axis) circuit input. The ratio between those measurements gives the ON/OFF ratio (top axis). Lastly, the time until 50% of the endpoint concentration of output protein is reached serves as a temporal measure of the circuit (t_50_; right axis). b) Endpoint concentrations, t_50_ and trade-offs for the CFFL with varying concentrations of σ^28^-encoding DNA construct. The highest ON/OFF ratio of 93x is reached for 0.5 nM σ^28^-encoding DNA construct. c) Trade-offs for varying concentrations of output DNA construct of the CFFL, where all ON/OFF ratios range between 4x and 7x. d) Trade-offs of the CFFL variants used in the composite CFFL (solid lines) and the trade-offs observed for both outputs of the circuit (dashed lines). The dashed green line represents the characteristics of the eGFP output with or without the input on the same side of the network (yellow shaded side), whereas the dashed blue line shows the characteristics of the eCFP output for its corresponding input trigger. All DNA constructs and concentrations are summarized in Table S3.

A wide range of σ^28^ expression levels was probed by varying the concentration of the DNA construct coding for the *E. coli* sigma factor (DNA **Y1**), whilst keeping other concentrations fixed. The CFFL was observed over time in the presence and absence of trigger DNA (DNA **X1**) to determine all its characteristics (Figure 3b). We observed a decrease in activation delay (*t_50_*) for increasing DNA **Y1** concentration, suggesting that σ^28^ expression is a key parameter to determine the dynamic behavior of the circuit. Endpoint expression exhibited an increase for higher DNA **Y1** concentrations, before plateauing and subsequently slightly decreasing. This behavior suggests that the σ^28^ protein concentration reaches a saturated regime and subsequent addition of more DNA merely limits the expression capacity available to the output protein, resulting in an inefficient use of resources and a decrease in circuit output. We observed similar behavior when varying the σ^28^ expression levels in the reference motif, except that the background expression decreased less compared to the CFFL when the σ^28^ expression strength was lowered (Figure S8). As a result, the CFFL motif displayed a higher *ON*/*OFF* ratio, reaching a value of 93x compared to 12x in the reference circuit.

Upon varying the concentration of output DNA species (DNA **Z1**), the delay in activation remained relatively constant, whilst the endpoint concentration and background endpoint concentration changed proportionally, with minimal changes to the *ON/OFF* ratio (Figure 3c). This further confirms that the σ^28^ DNA concentration is the main parameter influencing the temporal behavior of the CFFL and the DNA **Z1** concentration can merely be used to scale the circuit output. Moreover, the same intermediate DNA species dictates the fold-change of the circuit, since only when background expression from the σ^28^ DNA construct does not significantly activate the cognate promoter on the output DNA construct, can the overall leakage be minimized.

To investigate whether the same behavior persists in composite CFFL-based networks, we repeated the analysis on the composite CFFL with a shared intermediate node (Figure 3d). Since this network features two inputs and two outputs, the endpoint and transient characteristics were determined for each output in the presence and absence of the trigger that directly regulates the respective output (DNA **X2** for the eGFP output and DNA **X3** for the eCFP output; Figure 3d). We observed that the *ON/OFF* ratio of both outputs of the composite circuit is greatly reduced compared to CFFL 2 and 3, since the overall background expression of σ^28^ increased due to the presence of two σ^28^-producing constructs, which propagates into the background level of the output proteins. As a result, the background suppression behavior of the synthetic CFFL does not directly translate to the composite CFFL. Overall, our modular toehold switch-based CFFL system enabled the rapid analysis of a topologically complex synthetic genetic network.

### Time-varying circuit inputs

Characterization of the temporal behavior of CFFLs requires the introduction of input pulses of varying durations, which requires a method to dynamically add and eliminate DNA species in TXTL reactions. Here, we utilized semi-continuous microfluidic flow reactors^23,24,44^ in combination with TXTL reactions to implement and characterize the synthetic CFFL 1 and its reference motif, taking advantage of the controlled inflow and outflow capabilities of the reactors to implement variable length inputs (Figure 4a). After 3 or 4 hours of pre-equilibration without input DNA species, during which all constitutive and background expression equilibrated, persistent step inputs were applied to both the CFFL and reference circuits (Figure 4b). Like our analysis under batch conditions, endpoint and temporal characteristics were determined. We again observed that background expression in the absence of input is higher in the reference circuit, resulting in a larger *ON/OFF* ratio for the CFFL. The more efficient repression of background expression in the CFFL can be attributed to the sequential stages of repression achieved by the two toehold switches (Figure S9). Subsequently, DNA input pulses of lengths ranging from 15 minutes to 2 hours were applied to the CFFL to probe for noise-filtering behavior (Figure 4c). Square input pulses were emulated by initially supplying a high concentration of input DNA to create an immediate onset of signal, followed by a variable amount of regular concentration input steps to maintain a high input (Table S4). Finally, the input signal was terminated through omission of DNA **X1** from the inflow mixture, leading to a decrease of input trigger DNA concentration that was governed by the refresh rate of the microfluidic flow reactors (t_1/2_ = 25 min).

**Figure 4:**
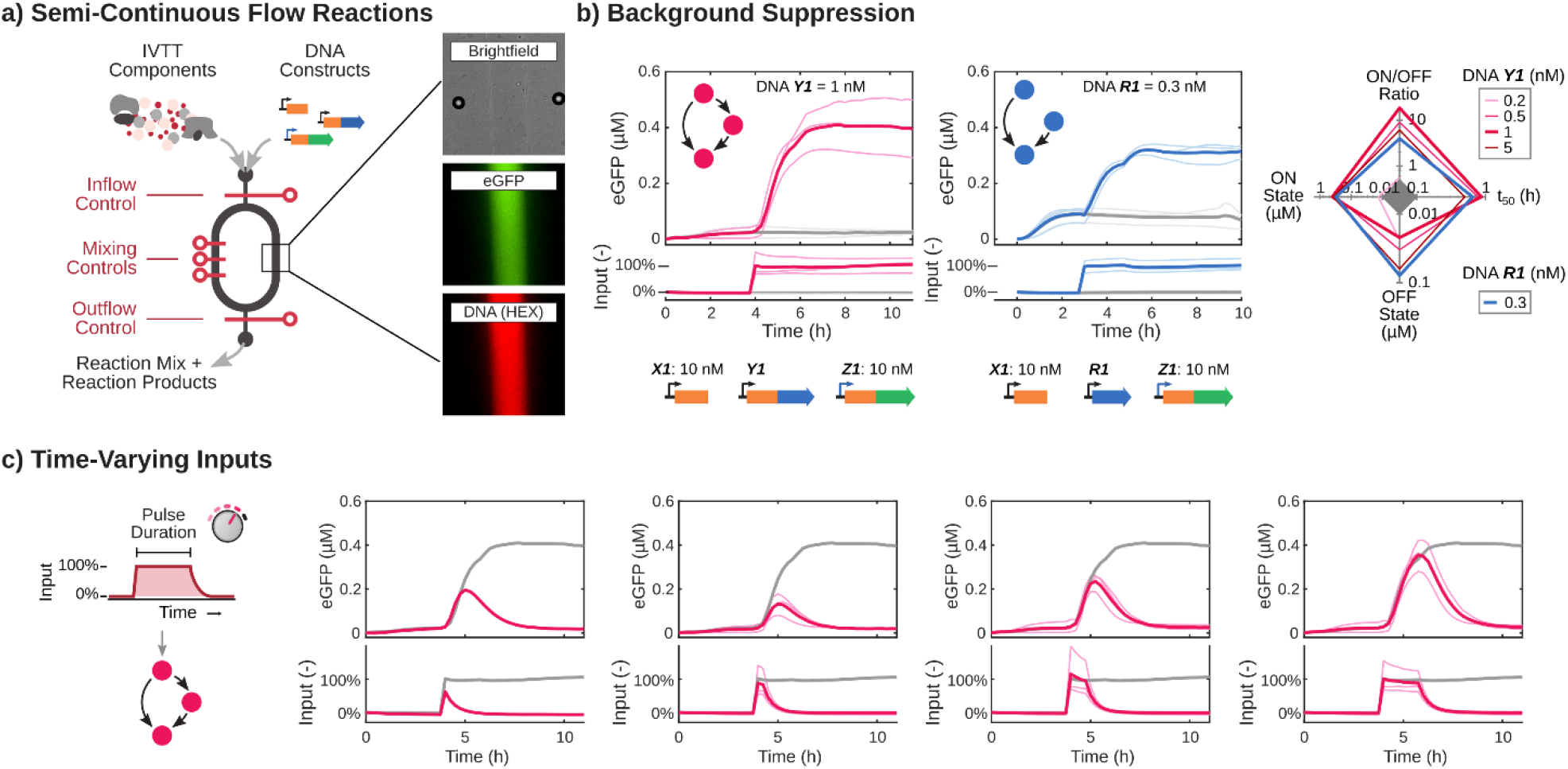
Semi-continuous flow reactions with time-varying inputs for the CFFL and reference circuits. a) Schematic drawing of a microfluidic semi-continuous flow reactor and its operation. In addition, an exemplar brightfield image and fluorescence micrographs of a channel in the reactor are shown. b) eGFP output time traces of flow reactions of the CFFL (left) and reference motif (right). Initially, no DNA encoding for the RNA input trigger was present. After either 3 or 4 hours, trigger DNA was added to the reactors to immediately reach a final concentration of 10 nM and was subsequently maintained at that concentration (colored lines). Negative controls, where no input DNA was added to the reactions are shown in gray. In addition, the trade-offs in characteristics of the flow reactions are plotted in a spider plot (CFFL in pink, reference motif in blue). The CFFL reaches a maximum ON/OFF ratio of 19x for 1 nM σ^28^-producing construct. c) Time-varying and corresponding CFFL circuit outputs (pink). A persistent input and corresponding output is shown in all plots as reference (gray). All DNA constructs and concentrations are summarized in Table S3 and the procedures to construct the time-dependent input pulses are described in Table S4.

We applied varying input pulse durations to the reference motif and CFFL, using multiple σ^28^ DNA concentrations, and determined the maximum GFP output for each input pulse as a measure of circuit response (Figure 5a). Short inputs elicited a response in both the CFFL and reference motif and we observed no clear indication of noise-filtering. Based on our observation that the σ^28^ DNA concentration is the main contributor to the dynamic behavior of the CFFL in batch TXTL reactions, we next examined the influence of DNA **Y1** concentration on the characteristics of CFFL 1 in semi-continuous flow reactions. While slightly different response dynamics were observed for varying σ^28^ expression strengths, short inputs still propagated through the CFFL motif. Taken together, these experimental results demonstrate that although the synthetic RNA-based CFFL does not display noise-filtering characteristics over a wide range of circuit parameters, it can be utilized to suppress background expression and yield a high fold-change.

**Figure 5:**
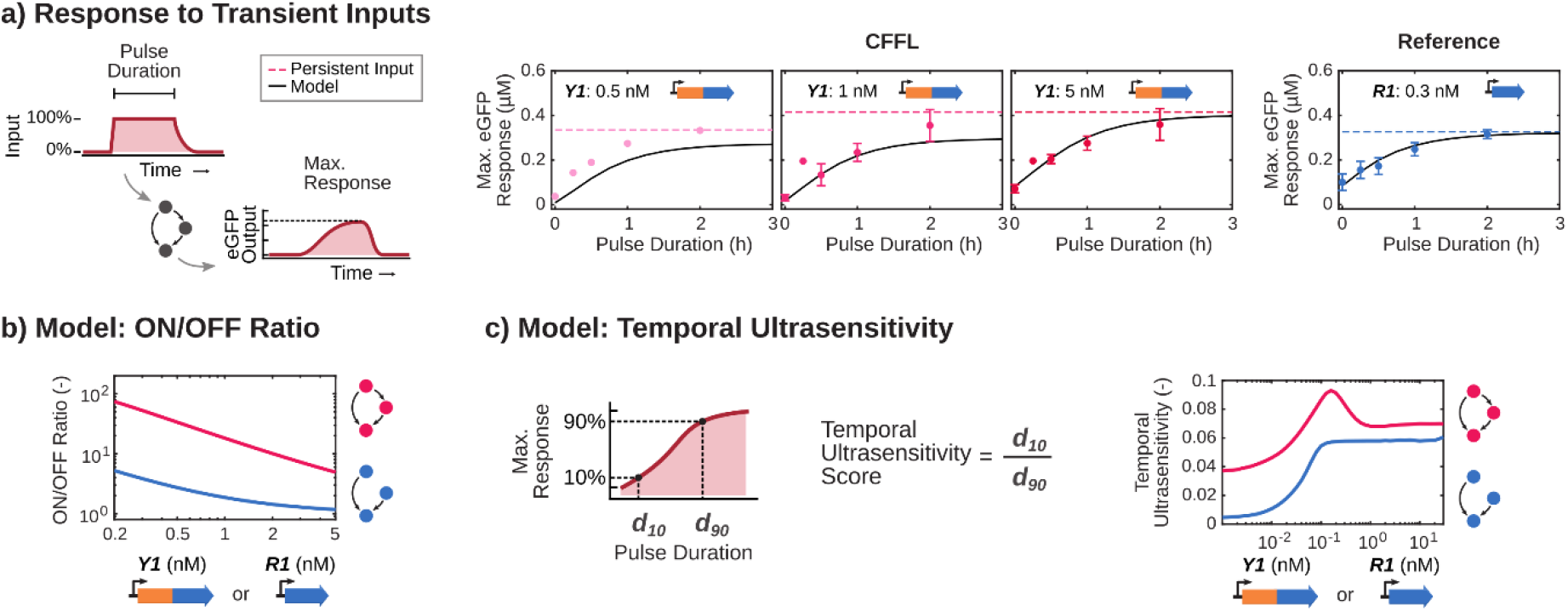
Computational analysis of the semi-continuous flow reactions with transient inputs. a) Maximum circuit responses for the CFFL (top plots) and reference motif (bottom plot) for no input, inputs of 15 min, 30 min, 1h, 2h (dots) and a constitutive input (dashed lines). The ODE model fit to this flow data (Equations S1-14; Table S2) is shown as black solid line. b) ODE model predictions for the ON/OFF ratio of the CFFL (pink) and reference motif (blue) under continuous flow conditions. The concentrations of the σ^28^-producing constructs were varied over the approximate range of experimental conditions and display comparable ON/OFF ratios to the experiments. c) ODE model predictions for the temporal ultrasensitivity of the CFFL (pink) and reference motif (blue) under continuous flow conditions. The concentration of the σ^28^-producing construct of each circuit was varied over a wide range of concentrations to explore the various behaviors that could be achieved using the circuits, but would be time-consuming to explore in vitro. All DNA constructs and concentrations are summarized in Table S3 and the procedures to construct the time-dependent input pulses are described in Table S4.

### Computational analysis of CFFL properties

To demonstrate that the observed experimental behaviors are general properties of the synthetic CFFL, we constructed ordinary differential equation (ODE) models of the CFFL and reference motif and parameterized the models using outputs of the flow reactor experiments and previously determined parameter values (Figure 5a, Supplementary Methods, Table S2).^25^ The models were analyzed to predict the *ON*/*OFF* ratio of the circuit output for varying concentrations of σ^28^ encoding DNA in both circuits (Figure 5b). The CFFL consistently produced a higher *ON*/*OFF* ratio than the reference circuit.

We further investigated the CFFL through the ODE model and determined temporal ultrasensitivities for varying concentrations of σ^28^ DNA (Figure 5c). The temporal ultrasensitivity measures how sharp the transition from 10% to 90% of the maximum output is with respect to the input pulse duration (d10 and d90, respectively; Equation S15).^35^ Temporal sensitivity quantifies filtering of short-lived inputs, since the transition from a low to high output for a change in input duration should be sharp for a noise filter. The synthetic CFFL displayed very low levels of temporal ultrasensitivity, which were only slightly higher than the reference motif, peaking in a narrow range of σ^28^ DNA concentrations around 0.2 nM.

We explored the temporal ultrasensitivity of the CFFL circuit further by modelling the circuit for a wide range of parameter values using Latin Hypercube sampling (Figure 6a; Table S2). The model displayed high temporal ultrasensitivity (>0.5) for only 0.3% of the parameter samples (Figure 6b). This observation is in line with a recent computational analysis, which revealed that the temporal ultrasensitivity of CFFL motifs has low robustness and is only significant in a small subset of circuit parameters.^35^ To investigate whether our synthetic CFFL could be improved to display stronger noise-filtering behavior, the sampled parameter space was filtered based on the computed temporal ultrasensitivity and statistics of the values of the selected parameter sets were determined. The parameter sets enriched for a high temporal ultrasensitivity were mainly associated with a high Hill-coefficient of the σ^28^ and DNA interaction (Figure 6c). Therefore, when incorporating a CFFL in synthetic genetic networks to increase tolerance to noise, a genetic regulator that binds more cooperatively to DNA should be utilized as delay element (Figure 6d). Alternatively, since cooperative transcription factors are scarce in prokaryotes,^46^ the sharpness of the σ^28^ binding curve could be increased using molecular titration with the anti-σ^28^ factor (FlgM) to achieve a similar effect.^47^ Nevertheless, adoption of our synthetic CFFL motif in synthetic circuits can prove to be beneficial in eliminating background expression and improving their fold-change.

**Figure 6:**
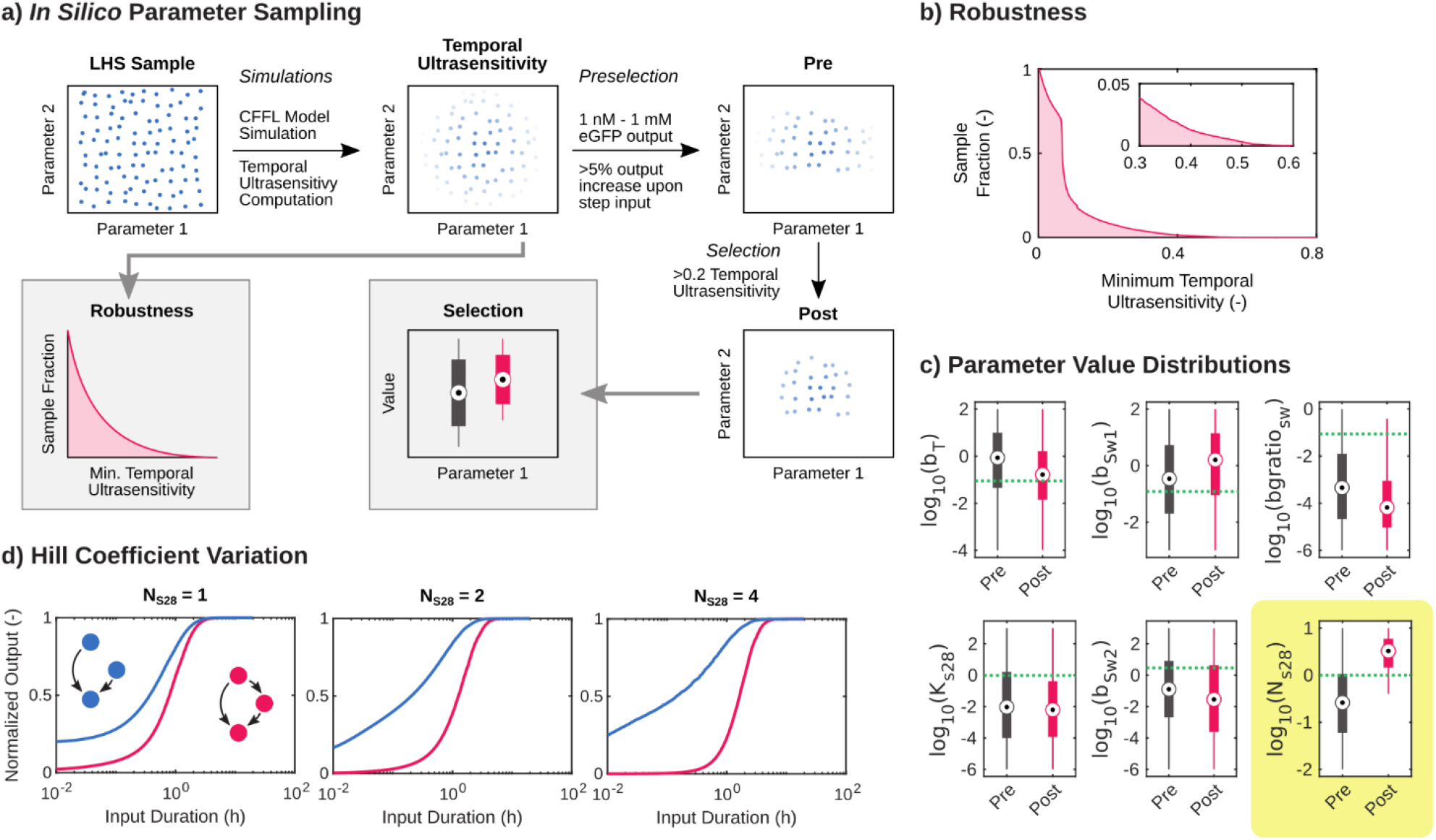
In silico parameter sampling of the CFFL ODE model. a) Schematic drawing of the sampling method and analysis procedures, shown for a 2-dimensional space for clarity. Latin hypercube sampling was used to create 10^5^ samples of the 13-dimensional logarithmic parameter space. For each parameter sample, the model was evaluated and the temporal ultrasensitivity was computed. To determine the robustness of the temporal ultrasensitivity behavior, for each temporal ultrasensitivity value the fraction of samples that displayed temporal ultrasensitivity of at least that value was computed. Additionally, the samples were filtered based on the computed properties. An initial selection was performed to create a collection of reasonable parameter values (pre). Subsequently, the samples were selected based on the computed temporal ultrasensitivity (post) These two collections were used to analyze parameter value distributions. b) The fraction of parameter samples that satisfied a minimum temporal ultrasensitivity threshold. A fraction of 0.3%, 8.5% and 21% displayed a temporal ultrasensitivity of at least 0.5, 0.2 and 0.1, respectively. c) The pre and post distributions of parameter values for a selection of parameters (see Figure S10 for all parameters and Table S2 for parameter descriptions and units). The dashed green lines show the parameter values used to simulate the experimental data in this research. A large increase in values between the pre and post sets can be observed for the σ^28^-binding Hill coefficient (N_S28_), which indicates that there is a preference for a high Hill coefficient when the temporal ultrasensitivity is high. d) Maximum output amplitudes of simulations of the CFFL and reference motif for a range of input pulse lengths, normalized to the maximum output for a step function input. Simulations with Hill coefficients 1, 2 and 4 demonstrate that the sharpness of the pulse length response of the CFFL increases for higher Hill coefficients, which is reflected in the associated temporal ultrasensitivity values of 0.09, 0.12 and 0.19, respectively.

## Conclusion

In this work, we reveal that toehold switch post-transcriptional regulators can be used to construct modular synthetic genetic networks in TXTL. We combined toehold switches with an *E. coli* sigma-factor to build a synthetic CFFL and a composite architecture based on naturally occurring organizations of feed-forward loops.^43^ The characteristics of the synthetic CFFL and composite CFFL were determined under batch TXTL conditions and in a semi-continuous microfluidic flow reactor. By comparing the circuit to a reference motif, we found that the synthetic CFFL could reduce background expression levels, thus increasing the fold-change of the circuit output. We utilized the flexibility of the microfluidic flow reactor to apply time-varying inputs to the synthetic CFFL, but could not identify temporal filtering in the circuit. *In silico* parameter sampling corroborated the observation that this behavior occurs for a small subset of circuit parameters.^35^

Since orthogonal alternative versions of the toehold switch^27,28^ and *E. coli* sigma factor^16,48^ are well characterized, we envision that all organizations of two CFFLs are suitable for implementation using our synthetic CFFL design, which enables characterization of topologically complex CFFL-based circuits that have as yet remained unexplored. Additionally, the modularity of the synthetic CFFL combined with the use of post-transcriptional regulation facilitates integration with existing genetic networks that are based on transcriptional control to provide repression of background expression to these networks. Whereas previous analyses of regulatory networks have focused on transcriptional interactions,^34^ recent work has shown that small RNAs (sRNA) play key roles in bacterial regulatory networks^26,30,49^ and sRNA-based feed-forward loops have been identified.^31,33,49,50^ As such, our CFFL-based circuits with RNA species as top regulators can provide a starting point for the characterization of these naturally occurring regulatory elements. Complemented by studies of the integration of synthetic genetic circuits into topologically more complex systems^51^ and the role of translational control in regulatory networks of cells,^30^ this work provides insight into the function of CFFLs and their application in synthetic biology.

## Methods

### Preparation of DNA Templates

DNA constructs were created with golden gate assembly (GGA) using the overlapping sequences adapted from Sun *et al.*^17^(Figure S1). The pBEST vector was a gift from Richard Murray and Vincent Noireaux (Addgene plasmid #45779) and was made suitable for GGA cloning using Gibson assembly (NEB Gibson Assembly Master Mix) of PCR products of the vector (NEB Phusion High-Fidelity DNA Polymerase) using primers pBEST_GA_1_F, pBEST_GA_1_R, pBEST_GA_2_F and pBEST_GA_2_R (Table S1). Promoters, UTR1, coding sequences and terminators were ordered from IDT as gBlock fragments or amplified from the pBEST vector using PCR. Toehold switch and trigger sequences were taken from previous studies in the group of Dr. P. Yin^27^ and PCR amplified. PCR products were gel purified using the QIAquick Gel Extraction Kit (Qiagen) and added in equimolar amounts to GGA assembly reactions with BSAI-HF (NEB), T4 ligase (Promega) and the T4 ligase buffer. GGA reactions were performed in a thermocycler according to a standard GGA protocol.^52^ The GGA products were transformed into NovaBlue cells (Merck), from which the plasmids were purified using the QIAprep Spin Miniprep Kit (Qiagen) and the DNA sequences were confirmed using Sanger sequencing.

Linear DNA templates for expression in IVTT reactions were created by PCR using Phusion High-Fidelity DNA Polymerase (NEB) with primers pBEST_LinL_F and pBEST_LinL_R (Table S1) and subsequent purification using QIAquick PCR Purification Kit (Qiagen).

### Preparation of Cell Lysate

The *E. coli* cell lysate was prepared from the RNAse E deficient BL21 STAR (DE3) (ThermoFisher Scientific) cells that were transformed with the pRARE vector from BL21 Rosettta (Merck). The lysate was prepared according to previously published protocols,^16,18^ with slight adaptations. The *E. coli* strain was grown in 2xYT medium supplemented with 40 mM potassium phosphate dibasic and 22 mM potassium phosphate monobasic until an OD600 of 1.7 was reached. The cultures were spun down and washed thoroughly with S30A buffer (14 mM Magnesium L-glutamate, 60 mM Potassium L-glutamate, 50 mM Tris, titrated to pH 8.2 using glacial acetic acid), before being resuspended in 0.9 mL S30A buffer per gram of dry pellet. The cell suspension was lysed using a French press at 16000 lb pressure in two passes and spun down. The supernatant was incubated at 37 °C for 1.5 hours and spun down. The supernatant was dialyzed into S30B buffer (14 mM Magnesium L-glutamate, 150 mM Potassium L-glutamate, titrated to pH 8.2 using 2 M Tris) in two steps for 3 hours total and spun down again. The supernatant was aliquotted, snap-frozen in liquid nitrogen and stored at −80 °C.

The energy mixture was prepared according to the protocol previously described by Sun *et al.*^18^ and a constant distribution amino acid solution was prepared.^53^

### Preparation of TXTL Reactions

The cell lysate (33% of total reaction volume) was combined with the energy mixture, amino acid solution (37.5 mM), Magnesium L-glutamate (8 mM), PEG-8000 (2%), GamS protein (3 μM; prepared as described by Sun *et al.*^17^) and MilliQ to form the 1.54x IVTT reaction mixture (65% of total reaction volume). The remaining volume (35%) of the reactions was used to add the linear DNA constructs of the gene networks and supplemented to the final volume with MilliQ. The DNA constructs and their concentrations in each experiment are summarized in Table S3.

### Batch TXTL Reactions

Batch IVTT reactions were prepared in total volumes of 10 μL and transferred to 384-wells Nunc plates. The reactions were incubated at 29 °C and eGFP and optionally eCFP fluorescence was measured on a Saffire II (Tecan), Spark 10M (Tecan) or Synergy H1M (Biotek) plate reader for at least 14 hours. The platereaders were calibrated using a titration range of purified eGFP protein. For the composite CFFL, where there are two outputs, eGFP and eCFP concentration ranges were measured in both the eGFP and eCFP measurement channels to determine the crosstalk between the two measurements.

### Microfluidic Device Fabrication and Flow TXTL Reactions

The microfluidic semi-continuous flow reactors were produced using standard soft lithography methods.^23,44^

Semi-continuous flow IVTT reactions were performed according to the protocol previously described by van der Linden *et al.*^44^, with adapted Labview control software to enable configuration of time-varying input signals. The 1.54x IVTT reaction mixture was stored on a water-cooled peltier element during the experiment to maintain reactivity of the solution, whereas other reactants were stored in tubing in the incubation chamber (29 °C). The input RNA trigger DNA template was mixed with a DNA-Hexachlorofluorescein conjugate (IDT; input_ref_hex, Table S1) to monitor and verify the applied circuit input pulses. Concentrations of the DNA constructs used are summarized in Table S3. Reactions were conducted for 11 h, during which 40% of each reactor was refreshed 15 minutes with 65% TXTL reaction mixture and 35% DNA or MilliQ. To create an input that resembles a square pulse function, an initial step containing 2.5x the final input concentration of DNA X1 was flushed in, after which all subsequent steps contained the regular input DNA concentration. The DNA solutions and operation sequences used to construct the time-dependent input pulses are described in Table S4. The devices were monitored on an Eclipse Ti-E inverted microscope (Nikon). Reactor channels were automatically detected in the obtained images using a custom Matlab (Mathworks) script and the average fluorescence of a 50×100 pixel rectangle at the center of a channel was calculated to represent the output fluorescence. After 11 hours the reactions were terminated and the microfluidic reactors were flushed with MilliQ. Optionally, microfluidic devices were cleaned for reuse through repeated flushing with a Terg-a-zyme enzyme detergent solution (Alconox).

### Parameter Fitting and Sampling

The ODE models of the CFFL and reference motif (Supplementary Methods, Equation S1-14) were implemented in Matlab (Mathworks) and numerically solved using the ode15s solver. We utilized the lsqnonlin solver using the trust-region-reflective algorithm to parameterize the ODE models. The model parameters were simultaneously fitted on a logarithmic scale to all flow reactor experimental data (Figure 4C), with 10^3^ Latin hypercube samples provided to the solver as initial parameter sets to prevent the fit from being only locally optimal. The fitted parameters and the resulting parameter set, that was used to perform further *in silico* experiments, is provided in Table S2.

To screen the behavior of the CFFL outside of the experimental parameter regime, Latin hypercube sampling (lhsdesign) was employed to generate 10^5^ parameter samples of wide range of parameter values in logarithmic space (see Table S2 for the parameters and their ranges). The CFFL ODE model was evaluated for all parameter samples to map the temporal ultrasensitivity of the CFFL. The network was simulated without input for 10 h, then 20 different time durations with input (0.01 – 20 evenly distributed on a logarithmic scale) and finally 4 h without input. From the maximum eGFP outputs of these 20 simulated experiments the temporal ultrasensitivity was computed (Equation S15). Parameter samples that resulted in a model that could not be correctly solved by ode15s were excluded from the analysis. To determine parameter values corresponding to a high temporal ultrasensitivity score, two stages of selection of the parameter samples were applied. First, samples were selected for a maximum eGFP output between 1 nM and 1 mM and a minimum increase of 5% of the maximum output upon addition of input trigger. The subsequent selection was conducted based on a minimum temporal ultrasensitivity of 0.2. The sampling and selection procedures are illustrated in Figure 6A and S10A.

## Acknowledgements

We thank Emilien Dubuc, Maaruthy Yelleswarapu and Roel Maas for helpful discussions. W.T.S.H. was supported by a TOPPUNT grant from The Netherlands Organization for Scientific Research (NWO). T.F.A.d.G. was supported by the NWO-VIDI grant from The Netherlands Organization for Scientific Research (NWO, 723.016.003). P.A.P., A.J.v.d.L. and T.F.A.d.G. were supported by an ERC starting grant by the European Research Council (project no. 677313 BioCircuit) and funding from the Ministry of Education, Culture and Science (Gravity programs, 024.001.035 and 024.003.013). J.K. was supported by the National Research Foundation of Korea (NRF-2019R1A2C1086830) grant funded by the Korean government (MSIT). P.Y. was supported by NSF CBET-1729397.

## References

(1) Alon, U. (2007) An Introduction to Systems Biology: Design Principles of Biological Circuits, 1st ed., Chapman &Hall/CRC, Boca Raton, FL..

(2) Davidson, E. H. (2006) The Regulatory Genome: Gene Regulatory Networks in Development and Evolution, Elsevier, Acad. Press,Amsterdam.

(3) Elowitz, M. B., and Leibler, S. (2000) A synthetic oscillatory network of transcriptional regulators. Nature 403, 335–338.

(4) Siuti, P., Yazbek, J., and Lu, T. K. (2013) Synthetic circuits integrating logic and memory in living cells. Nat. Biotechnol. 31, 448–452.

(5) Andrews, L. B., Nielsen, A. A. K., and Voigt, C. A. (2018) Cellular checkpoint control using programmable sequential logic. Science 361, eaap8987.

(6) Nissim, L., Wu, M.-R., Pery, E., Binder-Nissim, A., Suzuki, H. I., Stupp, D., Wehrspaun, C., Tabach, Y., Sharp, P. A., and Lu, T. K. (2017) Synthetic RNA-Based Immunomodulatory Gene Circuits for Cancer Immunotherapy. Cell 171, 1138–1150.e15.

(7) Roybal, K. T., Rupp, L. J., Morsut, L., Walker, W. J., McNally, K. A., Park, J. S., and Lim, W. A. (2016) Precision Tumor Recognition by T Cells With Combinatorial Antigen-Sensing Circuits. Cell 164, 770–779.

(8) Toda, S., Blauch, L. R., Tang, S. K. Y., Morsut, L., and Lim, W. A. (2018) Programming self-organizing multicellular structures with synthetic cell-cell signaling. Science 361, 156–162.

(9) Goentoro, L., Shoval, O., Kirschner, M. W., and Alon, U. (2009) The Incoherent Feedforward Loop Can Provide Fold-Change Detection in Gene Regulation. Mol. Cell 36, 894–899.

(10) Perez-Carrasco, R., Barnes, C. P., Schaerli, Y., Isalan, M., Briscoe, J., and Page, K. M. (2018) Combining a Toggle Switch and a Repressilator within the AC-DC Circuit Generates Distinct Dynamical Behaviors. Cell Syst. 6, 521–530.e3.

(11) Wall, M. E., Dunlop, M. J., and Hlavacek, W. S. (2005) Multiple Functions of a Feed-Forward-Loop Gene Circuit. J. Mol. Biol. 349, 501–514.

(12) Ma, W., Trusina, A., El-Samad, H., Lim, W. A., and Tang, C. (2009) Defining Network Topologies that Can Achieve Biochemical Adaptation. Cell 138, 760–773.

(13) Ahnert, S. E., and Fink, T. M. A. (2016) Form and function in gene regulatory networks: the structure of network motifs determines fundamental properties of their dynamical state space. J. R. Soc. Interface 13, 20160179.

(14) Payne, J. L., and Wagner, A. (2015) Function does not follow form in gene regulatory circuits. Sci. Rep. 5, 13015.

(15) Borkowski, O., Ceroni, F., Stan, G.-B., and Ellis, T. (2016) Overloaded and stressed: whole-cell considerations for bacterial synthetic biology. Curr. Opin. Microbiol. 33, 123–130.

(16) Garamella, J., Marshall, R., Rustad, M., and Noireaux, V. (2016) The All E. coli TX-TL Toolbox 2.0: A Platform for Cell-Free Synthetic Biology. ACS Synth. Biol. 5, 344–355.

(17) Sun, Z. Z., Yeung, E., Hayes, C. A., Noireaux, V., and Murray, R. M. (2014) Linear DNA for Rapid Prototyping of Synthetic Biological Circuits in an *Escherichia coli* Based TX-TL Cell-Free System. ACS Synth. Biol. 3, 387–397.

(18) Sun, Z. Z., Hayes, C. A., Shin, J., Caschera, F., Murray, R. M., and Noireaux, V. (2013) Protocols for Implementing an Escherichia coli Based TX-TL Cell-Free Expression System for Synthetic Biology. J. Vis. Exp. 50762, DOI: 10.3791/50762.

(19) Shin, J., and Noireaux, V. (2012) An E. coli Cell-Free Expression Toolbox: Application to Synthetic Gene Circuits and Artificial Cells. ACS Synth. Biol. 1, 29–41.

(20) Guo, S., and Murray, R. M. (2019) Construction of Incoherent Feedforward Loop Circuits in a Cell-Free System and in Cells. ACS Synth. Biol.8, 606–610.

(21) Patel, A., Murray, R. M., and Sen, S. (2020) Assessment of Robustness to Temperature in a Negative Feedback Loop and a Feedforward Loop. ACS Synth. Biol. 9, 1581–1590.

(22) Karzbrun, E., Tayar, A. M., Noireaux, V., and Bar-Ziv, R. H. (2014) Programmable on-chip DNA compartments as artificial cells. Science 345, 829–832.

(23) Niederholtmeyer, H., Stepanova, V., and Maerkl, S. J. (2013) Implementation of cell-free biological networks at steady state. Proc. Natl. Acad. Sci. 110, 15985–15990.

(24) Niederholtmeyer, H., Sun, Z. Z., Hori, Y., Yeung, E., Verpoorte, A., Murray, R. M., and Maerkl, S. J. (2015) Rapid cell-free forward engineering of novel genetic ring oscillators. eLife 4, DOI: 10.7554/eLife.09771.

(25) Yelleswarapu, M., van der Linden, A. J., van Sluijs, B., Pieters, P. A., Dubuc, E., de Greef, T. F. A., and Huck, W. T. S. (2018) Sigma Factor-Mediated Tuning of Bacterial Cell-Free Synthetic Genetic Oscillators. ACS Synth. Biol. 7, 2879–2887.

(26) Wagner, E. G. H., and Romby, P. (2015) Small RNAs in Bacteria and Archaea, in Advances in Genetics, pp 133–208. Elsevier, Amsterdam.

(27) Green, A. A., Silver, P. A., Collins, J. J., and Yin, P. (2014) Toehold Switches: De-Novo-Designed Regulators of Gene Expression. Cell 159, 925–939.

(28) Green, A. A., Kim, J., Ma, D., Silver, P. A., Collins, J. J., and Yin, P. (2017) Complex cellular logic computation using ribocomputing devices. Nature 548, 117–121.

(29) Lehr, F.-X., Hanst, M., Vogel, M., Kremer, J., Göringer, H. U., Suess, B., and Koeppl, H. (2019) Cell-Free Prototyping of AND-Logic Gates Based on Heterogeneous RNA Activators. ACS Synth. Biol. 8, 2163–2173.

(30) Nitzan, M., Rehani, R., and Margalit, H. (2017) Integration of Bacterial Small RNAs in Regulatory Networks. Annu. Rev. Biophys. 46, 131–148.

(31) Papenfort, K., Espinosa, E., Casadesús, J., and Vogel, J. (2015) Small RNA-based feedforward loop with AND-gate logic regulates extrachromosomal DNA transfer in *Salmonella*. Proc. Natl. Acad. Sci. 112, E4772–E4781.

(32) Tej, S., Gaurav, K., and Mukherji, S. (2019) Small RNA driven feed-forward loop: critical role of sRNA in noise filtering. Phys. Biol. 16, 046008.

(33) Thomason, M. K., Fontaine, F., De Lay, N., and Storz, G. (2012) A small RNA that regulates motility and biofilm formation in response to changes in nutrient availability in Escherichia coli: sRNA regulator of motility and biofilm formation. Mol. Microbiol. 84, 17–35.

(34) Milo, R. (2002) Network Motifs: Simple Building Blocks of Complex Networks. Science 298, 824–827.

(35) Gerardin, J., Reddy, N. R., and Lim, W. A. (2019) The Design Principles of Biochemical Timers: Circuits that Discriminate between Transient and Sustained Stimulation. Cell Syst. 9, 297–308.e2.

(36) Mangan, S., and Alon, U. (2003) Structure and function of the feed-forward loop network motif. Proc. Natl. Acad. Sci. 100, 11980–11985.

(37) Mangan, S., Zaslaver, A., and Alon, U. (2003) The Coherent Feedforward Loop Serves as a Sign-sensitive Delay Element in Transcription Networks. J. Mol. Biol. 334, 197–204.

(38) Purvis, J. E., Karhohs, K. W., Mock, C., Batchelor, E., Loewer, A., and Lahav, G. (2012) p53 Dynamics Control Cell Fate. Science 336, 1440–1444.

(39) Purvis, J. E., and Lahav, G. (2013) Encoding and Decoding Cellular Information through Signaling Dynamics. Cell 152, 945–956.

(40) Yosef, N., and Regev, A. (2011) Impulse Control: Temporal Dynamics in Gene Transcription. Cell 144, 886–896.

(41) Albeck, J. G., Mills, G. B., and Brugge, J. S. (2013) Frequency-Modulated Pulses of ERK Activity Transmit Quantitative Proliferation Signals. Mol. Cell 49, 249–261.

(42) Behar, M., Dohlman, H. G., and Elston, T. C. (2007) Kinetic insulation as an effective mechanism for achieving pathway specificity in intracellular signaling networks. Proc. Natl. Acad. Sci. 104, 16146–16151.

(43) Gorochowski, T. E., Grierson, C. S., and di Bernardo, M. (2018) Organization of feed-forward loop motifs reveals architectural principles in natural and engineered networks. Sci. Adv. 4, eaap9751.

(44) van der Linden, A. J., Yelleswarapu, M., Pieters, P. A., Swank, Z., Huck, W. T. S., Maerkl, S. J., and de Greef, T. F. A. (2019) A Multilayer Microfluidic Platform for the Conduction of Prolonged Cell-Free Gene Expression. J. Vis. Exp. e59655, DOI: 10.3791/59655.

(45) Stögbauer, T., Windhager, L., Zimmer, R., and Rädler, J. O. (2012) Experiment and mathematical modeling of gene expression dynamics in a cell-free system. Integr. Biol. 4, 494–501.

(46) Courey, A. J. (2001) Cooperativity in transcriptional control. Curr. Biol. 11, R250–R252.

(47) Buchler, N. E., and Louis, M. (2008) Molecular Titration and Ultrasensitivity in Regulatory Networks. J. Mol. Biol. 384, 1106–1119.

(48) Bervoets, I., Van Brempt, M., Van Nerom, K., Van Hove, B., Maertens, J., De Mey, M., and Charlier, D. (2018) A sigma factor toolbox for orthogonal gene expression in Escherichia coli. Nucleic Acids Res. 46, 2133–2144.

(49) Beisel, C. L., and Storz, G. (2011) The Base-Pairing RNA Spot 42 Participates in a Multioutput Feedforward Loop to Help Enact Catabolite Repression in Escherichia coli. Mol. Cell 41, 286–297.

(50) Mank, N. N., Berghoff, B. A., and Klug, G. (2013) A mixed incoherent feed-forward loop contributes to the regulation of bacterial photosynthesis genes. RNA Biol. 10, 347–352.

(51) Purnick, P. E. M., and Weiss, R. (2009) The second wave of synthetic biology: from modules to systems. Nat. Rev. Mol. Cell Biol. 10, 410–422.

(52) Engler, C., Kandzia, R., and Marillonnet, S. (2008) A One Pot, One Step, Precision Cloning Method with High Throughput Capability. PLoS ONE 3, e3647, DOI: 10.1371/journal.pone.0003647..

(53) Caschera, F., and Noireaux, V. (2015) Preparation of amino acid mixtures for cell-free expression systems. BioTechniques 58, 40–43.

