## Supporting Information for "Cell-Free Characterization of Coherent Feed-Forward Loop-Based Synthetic Genetic Circuits"

**Golden Gate Assembly**

```
plasmid...GCAT
          GCAT...promoter...AAGC            (variant with
                                            start codon)
                            AAGC...UTR...CATG
                                         CATG...CDS...TGAA
                            AAGC........trigger.......TGAA

                                            (variant 1 without
                                            start codon)
          GCAT...promoter.....switch.....GTTG
                                         GTTG...CDS...TGAA
                                            (variant 2 without
                                            start codon)
          GCAT...promoter.....switch.....CGAA
                                         CGAA...CDS...TGAA
                                                      TGAA...terminator...GTCG
                                                                          GTCG...plasmid
```

**Figure S1:** Schematic of the type of DNA components and their combinations to create the relevant DNA constructs.[1] The 5- to 3-prime 4 nucleotide Golden Gate Assembly (GGA) overhang sequences are shown in distinct colors, whereas the construct types (plasmid, promoter, UTR, switch, trigger, CDS, terminator) are displayed in black in between the overhangs. The start codon in the concerned overhang sequence is underlined. Two variants of the blue overhang sequence were used in combination with the toehold switches to decrease background expression by omitting the obsolete start codon in the original sequence.

| Primer | Sequence |
|---|---|
| pBEST_GA_1_F | GATACCGCGCGACCCACGCTCACC |
| pBEST_GA_1_R | GTGGGTCGCGCGGTATCATTGCAGCAC |
| pBEST_GA_2_F | CCTCAGGTAGGATGTAGGCCCTCAAAGAGATTGGCGCGGTGCTGG |
| pBEST_GA_2_R | GGCCTACATCCTACCTGAGGCGTTACCGGACCAGAAGTTGTCCTGGC |
| pBEST_LinL_F | GGCGAATCCTCTGACCAGC |
| pBEST_LinL_R | CCAAGCTGGACTGTATGCACG |
| input_hex_ref | /5HEX/GGCGAATCCTCTGACCAGC |
| pPBEST_GGA_OL5_F | ATATAGGTCTCTGTCGCGGCATGATAAGCTGTCAAACATG |
| pPBEST_GGA_OL1_R | GTCCTGGTCTCTATGCGTTACCGGACCAGAAGTTGTC |
| P70a_GGA_OL1_F | AGAACGGTCTCAGCATGATGTCTGAGCTAACACCGTGCGTG |
| P70a_GGA_OL2_R | AGCCAGGTCTCAGCTTTGCAACCATTATCACCGCCAG |
| P28a_GGA_OL1_F | AGAACGGTCTCAGCATGATGTCCAGGACAACTTCTGGTCCGG |
| P28a_GGA_OL2_R | AGCCAGGTCTCAGCTTTTGATCTCGTTATCGGCAAGGAG |
| TriggerA_GGA_OL2_F | AGCCAGGTCTCAAAGCGGGATACACATAGAATCATGTGTATAAC |
| TriggerA_GGA_OL4_R | TTAGTGGTCTCATTCAGCTAGTGATTGAATATGATAGAAGTTTAGTAG |
| TriggerB_GGA_OL2_F | AGCCAGGTCTCAAAGCGGGACCGCAATGCGGAAATTG |
| TriggerB_GGA_OL4_R | TTAGTGGTCTCATTCACAAGGGGTTATGCTATTCGCCTCTATTC |
| S28_GGA_OL3_F | AACAGGGTCTCACATGAATTCACTCTATACCGCTGAAGG |
| S28_GGA_OL3S_F | AACAGGGTCTCACATGAATTCACTCTATACCGCTGAAGG |
| S28_GGA_OL4_R | TCCCCGGTCTCATTCATAACTTACCCAGTTTAGTGCGTAACC |
| e(G/C)FP_OL3S_F | AACAGGGTCTCAGTTGGTGAGCAAGGGCGAGG |
| e(G/C)FP_OL3SS_F | AACAGGGTCTCACGAAGAGCTTTTCACTGGCGTTGTTCC |
| e(G/C)FP_OL4_R | TTAGTGGTCTCATTCACTTGTACAGCTCGTCCATGC |

**Table S1:** Primer sequences used in this research. GGA = Golden Gate Assembly, GA = Gibson Assembly and OL***X*** represents for overlap sequence ***X*** (see Figure S1 for the various overlap sequences).

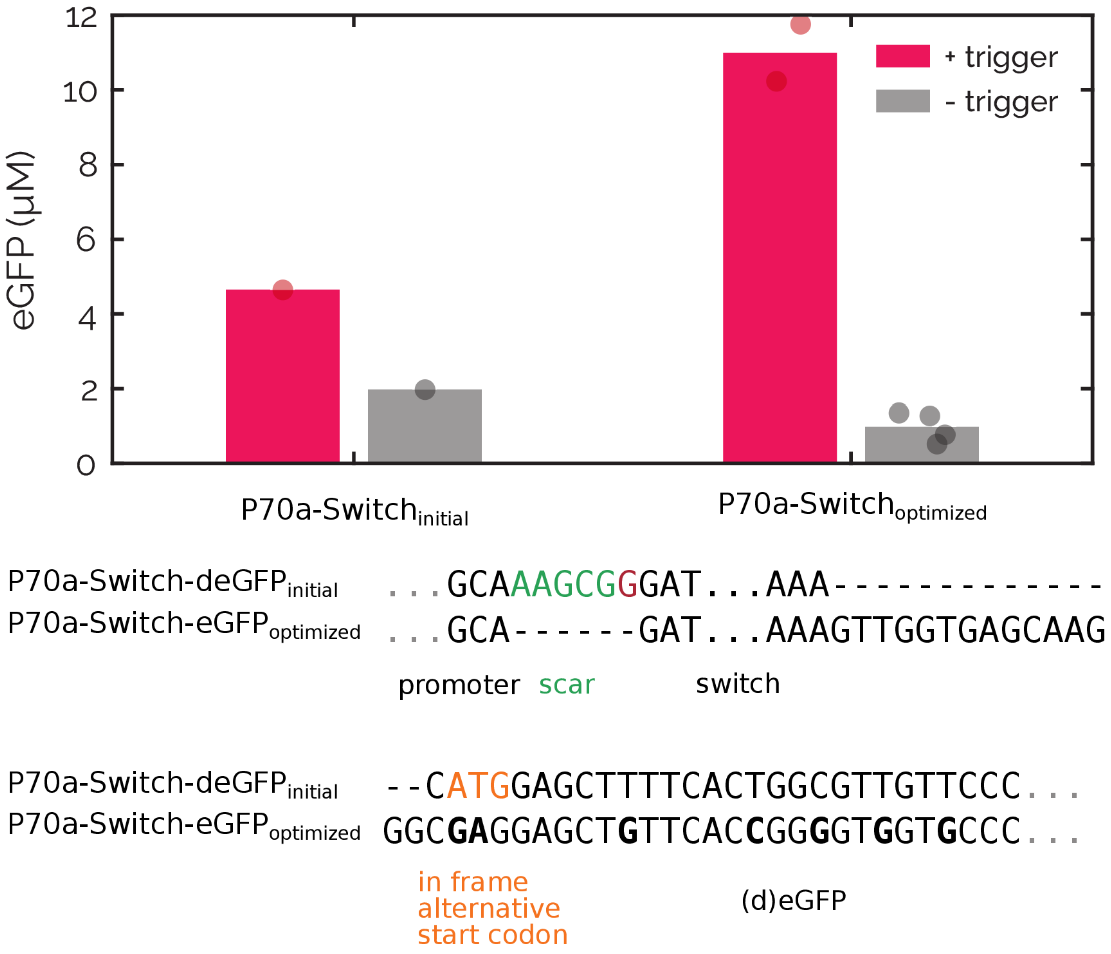


**Figure S2:** Comparison of the expression levels and sequence of an initial toehold switch-bearing DNA construct design and an optimized version. The differences in sequence at the promoter-switch transition and switch-CDS transition are displayed using sequence alignment at the aforementioned positions. Deletions are shown as a dash (-) and mutations are displayed in bold. The most notable changes are the deletion of a cloning scar between the promoter and switch and the omission of the start codon at the start of the eGFP sequence, which only leaves the toehold switch-regulated start codon for initiation of translation. The expression levels with (pink) and without (gray) toehold trigger encoding DNA construct are shown in the graph. The bars represent the average expression after 14 hours and the points represent individual experiments. All experimental conditions are summarized in Table S3.

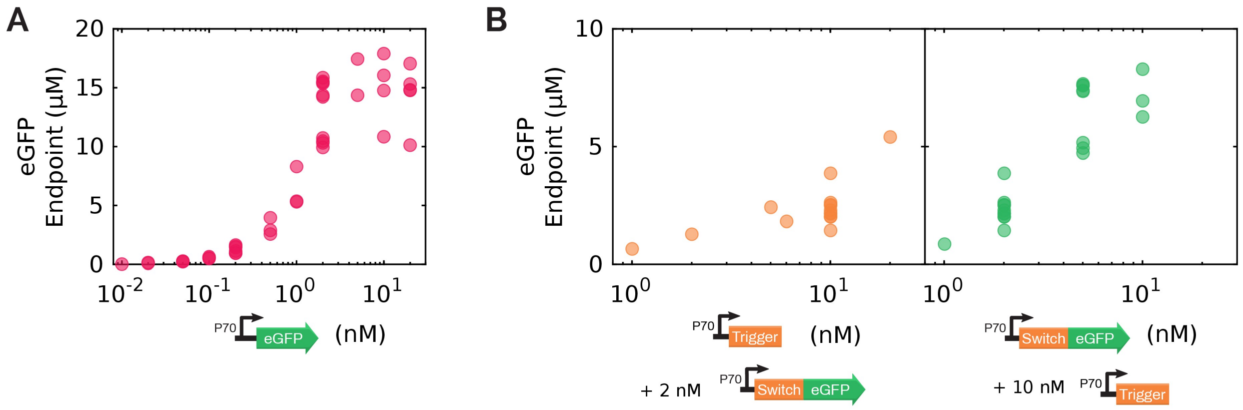


**Figure S3:** A) Endpoint eGFP output concentrations of a titration of the P70a-eGFP DNA construct, which functions as an expression positive control construct. Expression from this construct plateaus for DNA concentrations above ~5 nM. B) Endpoint eGFP concentrations for a titration of trigger (left) and switch DNA constructs (right). Both experiments display an increase in eGFP output for increasing concentrations of the DNA constructs, reaching expression levels close to the positive control (A). All experimental conditions are summarized in Table S3.

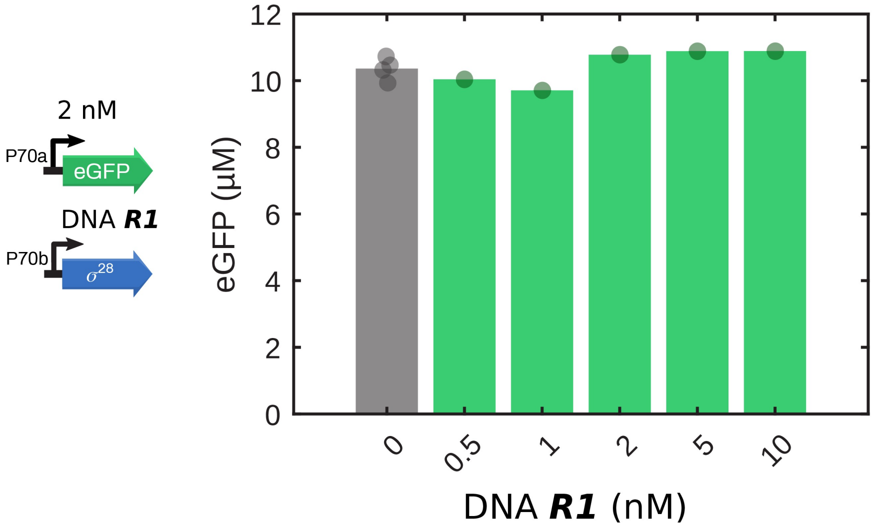


**Figure S4:** Endpoint expression of a simple eGFP gene in the presence of various concentrations of $\sigma^{28}$-producing DNA construct. The $\sigma^{28}$ protein produced by the DNA ***R1*** construct competes with $\sigma^{70}$ for the RNA polymerase. Nevertheless, the production of eGFP from the $\sigma^{70}$-controlled P70 promoter did not decrease for increasing concentrations of DNA ***R1***, indicating that the concentration of $\sigma^{28}$ protein that is produced is too low to significantly alter $\sigma^{70}$-regulated protein expression. All experimental conditions are summarized in Table S3.

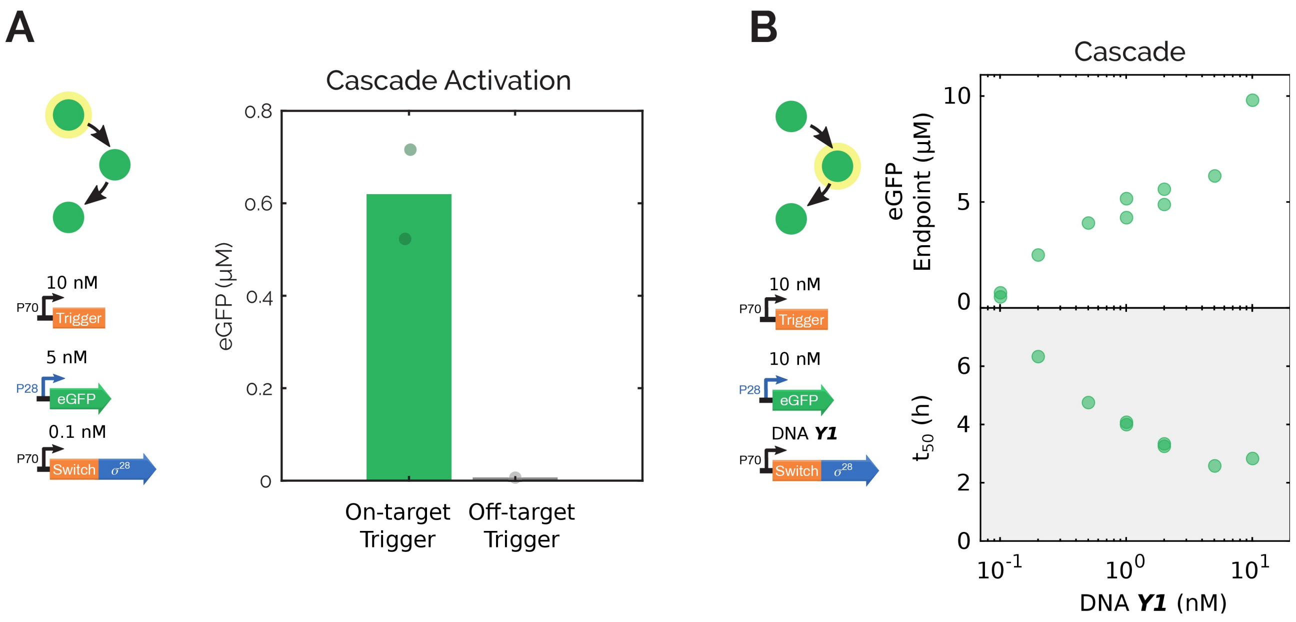


**Figure S5:** A) Endpoint expression of the expression cascade consisting of a toehold trigger that activates a toehold switch that regulates the translation of $\sigma^{28}$, which activates the transcription of an eGFP output gene. The cascade produced eGFP when an on-target toehold trigger was used, whilst expression was absent when an off-target trigger was used instead. B) The endpoint eGFP concentration and expression delay ($t_{50}$) for a range of $\sigma^{28}$ construct concentrations. Expression levels increased for increasing $\sigma^{28}$ DNA concentrations, while the $t_{50}$ steadily dropped. All experimental conditions are summarized in Table S3.

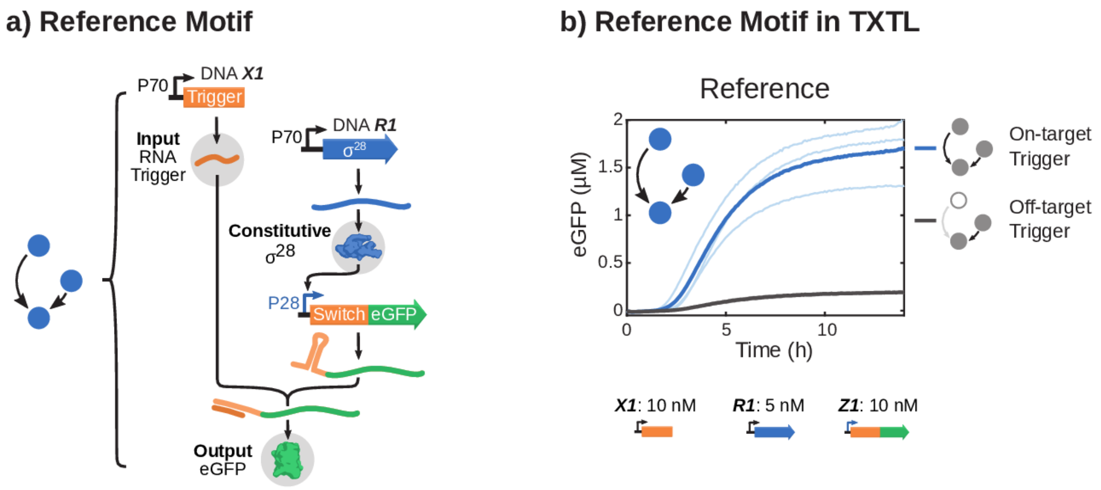


**Figure S6:** a) Schematic representation of all DNA species and the DNA, RNA and protein level interactions that constitute the reference motif. The three main components are the toehold switch, marked by the 5'-adjacent RNA stem loop, and its corresponding RNA trigger (orange), the E. coli $\sigma^{28}$-factor (blue) and fluorescent output protein (green). b) Time traces of eGFP expression of the reference motif with an on-target trigger (light blue traces are three distinct experiment and dark blue is their average) and an off-target trigger (dark gray). All experimental conditions are summarized in Table S3.

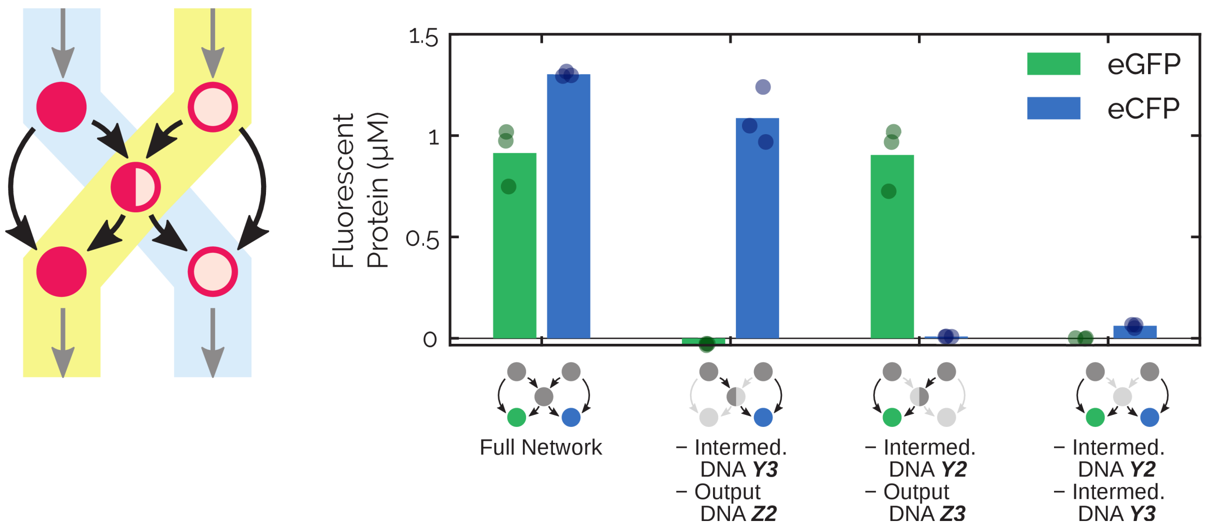


**Figure S7:** Endpoint eGFP (green) and eCFP (blue) concentrations of select subnetworks of the composite CFFL, where only the trigger and $\sigma^{28}$ constructs from one side of the network and the trigger and output constructs from the other side were expressed. These experiments demonstrate that the two sides of the network can influence each other through the $\sigma^{28}$ factor. The full composite CFFL and the composite CFFL where all $\sigma^{28}$ constructs are left out are included as positive and negative control, respectively. Bars represent the average output protein endpoint concentration and the data points display the endpoint concentration of individual experiments. DNA concentrations are, unless omitted in the subnetwork, as shown in Figure 2B. All experimental conditions are summarized in Table S3.

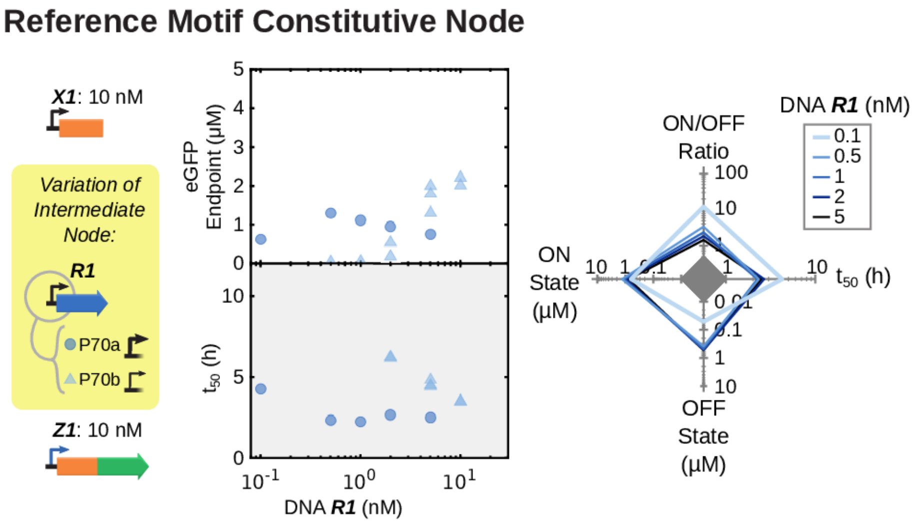


**Figure S8:** Endpoint concentrations, $t_{50}$ and trade-offs for the reference motif with varying concentrations of $\sigma^{28}$-encoding DNA construct (DNA R1). In addition to the default construct (circles), a construct with a point mutation in the promoter to give a low efficiency $\sigma^{28}$-producing gene was evaluated (triangles). The trade-offs are shown for the default high-yield $\sigma^{28}$ DNA construct. All experimental conditions are summarized in Table S3.

### Ordinary differential equation (ODE) models

We constructed ODE models of the synthetic CFFL and reference motif to simulate flow reactor experiments. Protein species and all RNA species and complexes were taken as state variables of the ODE models. DNA species were defined as parameters that can be adjusted to emulate the dynamic inputs given in the flow experiments. Complex formation of the RNA toehold switches and triggers, constitutive transcription and translation were modeled with a mass action term, while activation of the P28 promoter by the $\sigma^{28}$ factor was modeled using Hill-type kinetics. Fluorescent protein maturation was not explicitly included, but is captured in the translation rate of the protein. Since the model was exclusively used to model flow reactor experiments, depletion of resources was assumed not to play a major role and was not represented in the ODE models.

#### *ODE Model of the synthetic CFFL*

The CFFL model comprises a RNA trigger (*T*) species that is produced from DNA species ***X1*** ($DNA_T$) with rate $b_T$. SwitchA-S28 mRNA is produced from DNA species ***Y1*** ($DNA_{Sw1}$) with rate $b_{Sw1}$. SwitchA-eGFP mRNA is produced by expression from DNA species ***Z1*** ($DNA_{Sw2}$) with rate $B_{Sw2}$ (background expression) and with rate $b_{Sw2}$ upon activation by the $\sigma^{28}$ factor (*S28*) through Hill kinetics ($K_{S28}$ and $N_{S28}$). The $\sigma^{28}$ factor was previously shown to behave according to Michaelis-Menten kinetics,[2] so $N_{S28}$ was set to 1 for the parameter fitting procedure, but was varied during parameter sampling (Table S2). Binding of trigger RNA (*T*) to SwitchA-S28 mRNA (*Sw1*) and SwitchA-eGFP mRNA (*Sw2*) to form complexes *TSw1* and *TSw2*, respectively, was explicitly modeled using mass action kinetics ($k_{on,TSw}$ and $k_{off,TSw}$). Translation from the trigger:switch complexes was represented by rates $k_{tl,S28}$ and $k_{tl,GFP}$. Since translation from unbound switch RNA species can also occur, but at a lower rate determined by the efficiency of the toehold switch, this was included in the model using $bgratio_{sw}$. The resulting translation rate of unbound SwitchA-S28 mRNA is the product of $k_{tl,S28}$ and $bgratio_{sw}$ and the translation rate of unbound SwitchA-eGFP mRNA is the product of $k_{tl,GFP}$ and $bgratio_{sw}$. The degradation rates of all RNA species was represented by parameter $a_{RNA}$ and of all protein species by parameter $a_{protein}$. Since the degradation rate of proteins in the TXTL reactions was negligible, as demonstrated by the constant plateaus after 10+ hours of batch TXTL reactions, the value of $a_{protein}$ was set to zero. The flow rate of the experiment can be found in all equations as parameter $k_{flow}$, which was set as 1.6 $h^{-1}$ for all experiments, giving a residence time of 37.5 min. The resulting ODE model of the synthetic CFFL is given by Equations S1-8, where Equation S5 shows a helper function *f(..)* which is used in the Hill equation in Equation S6.

$$\frac{dT(t)}{dt} = b_T \cdot DNA_T + k_{off,TSw} \cdot TSw1(t) - k_{on,TSw} \cdot T(t) \cdot Sw1(t) + k_{off,TSw} \cdot TSw2(t) - k_{on,TSw} \cdot T(t) \cdot Sw2(t) - (a_{RNA} + k_{flow}) \cdot T(t) \quad \text{(S1)}$$

$$\frac{dSw1(t)}{dt} = b_{Sw1} \cdot DNA_{Sw1} + k_{off,TSw} \cdot TSw1(t) - k_{on,TSw} \cdot T(t) \cdot Sw1(t) - (a_{RNA} + k_{flow}) \cdot Sw1(t) \quad \text{(S2)}$$

$$\frac{dTSw1(t)}{dt} = k_{on,TSw} \cdot T(t) \cdot Sw1(t) - k_{off,TSw} \cdot TSw1(t) - (a_{RNA} + k_{flow}) \cdot TSw1(t) \quad \text{(S3)}$$

$$\frac{dS28(t)}{dt} = k_{tl,S28} * TSw1(t) + k_{tl,S28} \cdot bgratio_{sw} \cdot Sw1(t) - (a_{protein} + k_{flow}) \cdot S28(t) \quad \text{(S4)}$$

$$f(S28(t)) = \left(\frac{S28(t)}{K_{S28}}\right)^{N_{S28}} \quad \text{(S5)}$$

$$\frac{dSw2(t)}{dt} = B_{Sw2} \cdot DNA_{Sw2} + b_{Sw2} \cdot DNA_{Sw2} \cdot \frac{f(S28(t))}{1 + f(S28(t))} + k_{off,TSw} \cdot TSw2(t) - k_{on,TSw} \cdot T(t) \cdot Sw2(t) - (a_{RNA} + k_{flow}) \cdot Sw2(t) \quad \text{(S6)}$$

$$\frac{dTSw2(t)}{dt} = k_{on,TSw} \cdot T(t) \cdot Sw2(t) - k_{off,TSw} \cdot TSw2(t) - (a_{RNA} + k_{flow}) \cdot TSw2(t) \quad \text{(S7)}$$

$$\frac{dGFP(t)}{dt} = k_{tl,GFP} \cdot TSw2(t) + k_{tl,GFP} \cdot bgratio_{sw} \cdot Sw2(t) - (a_{protein} + k_{flow}) \cdot GFP(t) \quad \text{(S8)}$$

Where species *T* is the trigger RNA, *Sw1* the Switch-S28 RNA and *TSw1* the complex of bound Trigger and Switch-S28 RNA. *Sw2* is the Switch-eGFP RNA and *TSw2* the complex of Trigger and Switch-eGFP RNA. *S28* is the $\sigma^{28}$ protein species and *GFP* is the eGFP protein species. $DNA_X$ is the concentrations of the DNA coding for RNA species **X** (nM). Further parameter descriptions and units can be found in Table S2.

##### *ODE Model of the reference motif*

The model of the reference motif is largely equal to the CFFL model, except for the absense of an interaction between RNA trigger *T* and mRNA species Sw1. In the reference motif model, RNA species Sw1 represents the S28 RNA (without toehold switch) that results from transcription of DNA species ***R1*** ($DNA_{Sw1}$). Since parameter $DNA_{Sw1}$ now represents the concentration of a different physical DNA species (***R1*** instead of ***Y1***), the transcription rate of this species ($b_{Sw1,ref}$) is assumed to be different than for the CFFL model.

$$\frac{dT(t)}{dt} = b_T \cdot DNA_T + k_{off,TSw} \cdot TSw2(t) - k_{on,TSw} \cdot T(t) \cdot Sw2(t) - (a_{RNA} + k_{flow}) \cdot T(t) \quad \text{(S9)}$$

$$\frac{dSw1(t)}{dt} = b_{Sw1,ref} \cdot DNA_{Sw1} - (a_{RNA} + k_{flow}) \cdot Sw1(t) \quad \text{(S10)}$$

$$\frac{dS28(t)}{dt} = k_{tl,S28} * Sw1(t) - (a_{protein} + k_{flow}) \cdot S28(t) \quad \text{(S11)}$$

$$\frac{dSw2(t)}{dt} = B_{Sw2} \cdot DNA_{Sw2} + b_{Sw2} \cdot DNA_{Sw2} \cdot \frac{f(S28(t))}{1+f(S28(t))} + k_{off,TSw} \cdot TSw2(t) - k_{on,TSw} \cdot T(t) \cdot Sw2(t) - (a_{RNA} + k_{flow}) \cdot Sw2(t) \quad \text{(S12)}$$

$$\frac{dTSw2(t)}{dt} = k_{on,TSw} \cdot T(t) \cdot Sw2(t) - k_{off,TSw} \cdot TSw2(t) - (a_{RNA} + k_{flow}) \cdot TSw2(t) \quad \text{(S13)}$$

$$\frac{dGFP(t)}{dt} = k_{tl,GFP} \cdot TSw2(t) + k_{tl,GFP} \cdot bgratio_{sw} \cdot Sw2(t) - (a_{protein} + k_{flow}) \cdot GFP(t) \quad \text{(S14)}$$

Where species *T* is the trigger RNA and *Sw1* is the S28 RNA (without any toehold switch). *Sw2* is the Switch-eGFP RNA and *TSw2* the complex of Trigger and Switch-eGFP RNA. *S28* is the $\sigma^{28}$ protein species and *GFP* is the eGFP protein species. $DNA_X$ is the concentrations of the DNA coding for RNA species **X** (nM). Further parameter descriptions and units can be found in Table S2.

### Temporal Ultrasensitivity Computation

The temporal ultrasensitivity calculations were based on the α and β measures, calculated from the maximum output amplitude per input pulse length (Figure 4C and D).[3] α is the pulse duration at which the maximum response has increased 10% of the total difference between background expression (no input) and full activation (persistent step input), computed by linearly interpolating the nearest experimental or simulation data points. Similarly, β marks the pulse duration of 90% increase in response. The temporal ultrasensitivity is defined as follows:

$$temporal\,ultrasensitivity=\frac{\alpha}{\beta} \tag{S15}$$

Thus, the temporal ultrasensitivity is low for a gradually increasing activation upon an increase in input pulse length and approaches 1 for a circuit that approaches immediate switch-like behavior at a given pulse length.

| Parameter Name | Unit | Description | Value | Sampling bounds |
|---|---|---|---|---|
| $k_{flow}$ | $h^{-1}$ | Microfluidic reactor flow rate | 1.6 | - |
| $k_{on,TSw}$ | $\mu M^{-1}h^{-1}$ | Trigger and switch association rate | $^{0}$ | $^{0}$ |
| $k_{off,TSw}$ | $h^{-1}$ | Trigger and switch dissociation rate | $^{0}$ | $10^{-12}$ - $10^{-3}$ |
| $b_T$ | $\mu M\ h^{-1}$ $(nM\ DNA)^{-1}$ | Trigger transcription rate | $0.09^*$ | $^{0}$ |
| $a_{RNA}$ | $h^{-1}$ | RNA degradation rate | $7.9^*$ | $^{0}$ |
| $b_{Sw1}$ | $\mu M\ h^{-1}$ $(nM\ DNA)^{-1}$ | Switch-S28 construct transcription rate | $0.12^*$ | $^{0}$ |
| $k_{tl,S28}$ | $h^{-1}$ | S28 translation rate | 15 | $^{0}$ |
| $bgratio_{Sw}$ | - | Fraction of translation rate observed from unbound switch | $0.09^*$ | $^{0}$ |
| $a_{protein}$ | $h^{-1}$ | Protein degradation rate | 0 | - |
| $K_{S28}$ | $\mu M$ | S28 binding constant | $0.96^*$ | $^{0}$ |
| $N_{S28}$ | - | Hill coefficient | 1 | $^{0}$ |
| $B_{Sw2}$ | $\mu M\ h^{-1}$ $(nM\ DNA)^{-1}$ | Background expression of Switch-eGFP construct. | 0 | - |
| $b_{Sw2}$ | $\mu M\ h^{-1}$ $(nM\ DNA)^{-1}$ | Switch-eGFP construct transcription rate | $2.88^*$ | $^{0}$ |
| $k_{tl,GFP}$ | $h^{-1}$ | eGFP translation rate | 4.5 | $^{0}$ |
| $b_{Sw1,ref}$ | $\mu M\ h^{-1}$ $(nM\ DNA)^{-1}$ | S28 reference construct transcription rate | $0.18^*$ | $^{0}$ |
| $DNA_T$ | nM | Trigger DNA (***X1***) concentration. | 10 | - |
| $DNA_{Sw1}$ | nM | DNA ***Y1*** (CFFL) or ***R1*** (Reference) concentration. | 1 | - |
| $DNA_{Sw2}$ | nM | Output DNA (***Z1***) concentration. | 10 | - |

**Table S2:** Model parameter, their units, used values and sampled parameter range. Sample values marked with an asterisk ($^*$) were obtained through parameter fitting. The DNA concentrations

($DNA_T$, $DNA_{Sw1}$ and $DNA_{Sw2}$) are the concentrations used for parameter sampling. The concentrations used in other *in silico* experiments are given in Table S3.

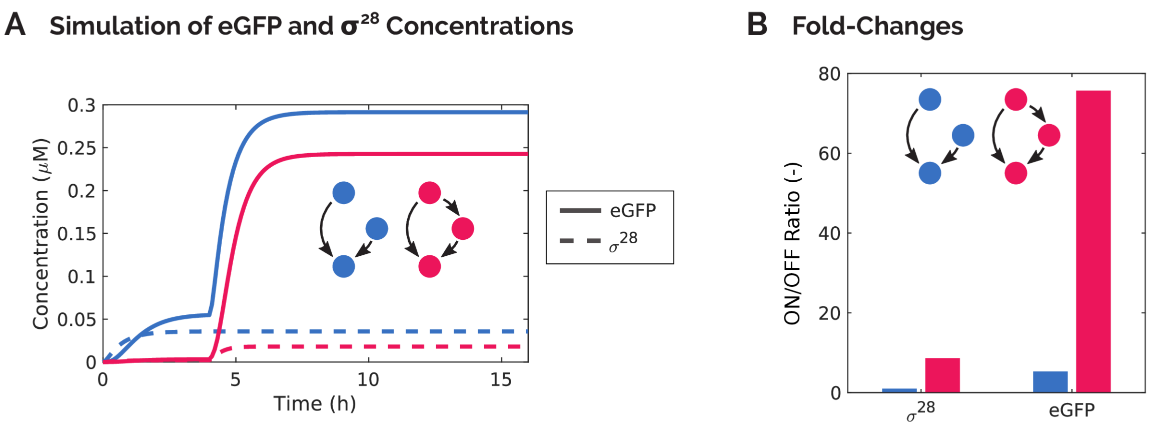


**Figure S9:** Simulations of the influence of the presence of toehold switches at two positions in the CFFL circuit. A) Simulated concentration traces for the CFFL (pink) and reference motif (blue) of both the eGFP output (solid) and $\sigma^{28}$ protein concentration. B) The *ON/OFF* ratios of $\sigma^{28}$ and eGFP in the CFFL and reference circuits. The $\sigma^{28}$ concentration of the reference motif remains constant after addition of an input, since it is constitutively produced. The eGFP output of the reference motif displays an *ON/OFF* ratio of 8.6, which is caused only by the toehold switch on the output construct, since the other switch is not present in the reference motif. The fold-change in the $\sigma^{28}$ concentration of the CFFL mainly represents the activation of the toehold switch on the switch-$\sigma^{28}$ construct and was observed to be 5.3. Together, the activation of the switches largely accounts for the high fold-change in eGFP output of the CFFL, leaving a factor 1.7 to be accounted for by other factors, such as differences in parameter values between the reference motif and CFFL and non-linearity of the $\sigma^{28}$-DNA interaction.

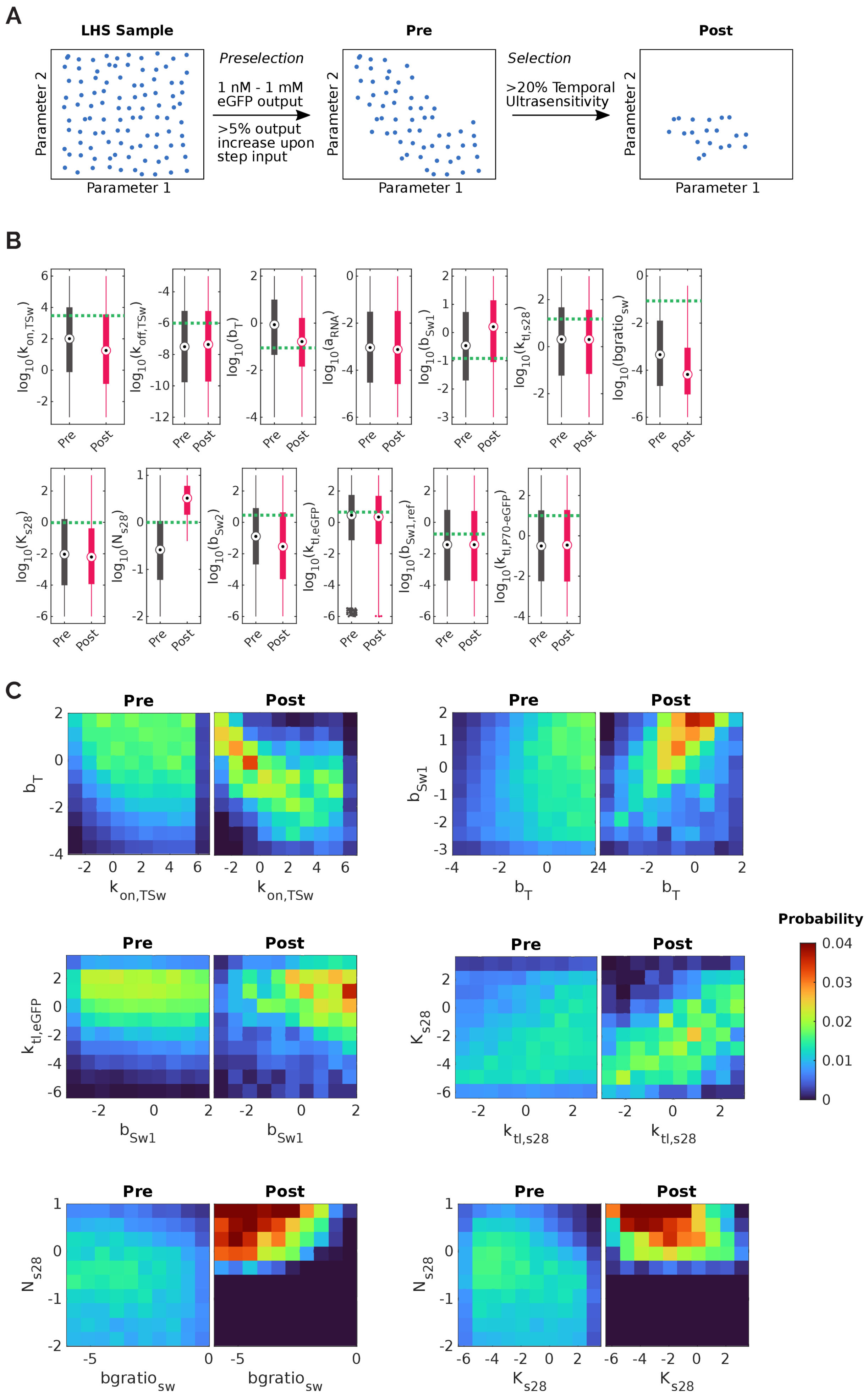

A
LHS Sample
Pre
Post
Preselection
1 nM - 1 mM eGFP output
>5% output increase upon step input
Selection
>20% Temporal Ultrasensitivity
Parameter 1
Parameter 2
B
Pre
Post
C
Pre
Post
Probability

**Figure S10:** *In silico* selection of parameter samples of the CFFL model that display high temporal ultrasensitivity. A) Schematic that shows the selection procedure of the parameter samples. First, a latin hypercube sampling (LHS) of the 13-dimensional parameter space was taken in the logarithmic domain. For each parameter sample, the temporal ultrasensitivity and general output statistics (maximum output and relative activation upon when given a step input) of the CFFL were computed. An initial selection was performed based on the general output measures to ensure reasonable output protein concentrations (between 1 nM and 1 mM) and discernible activation when an input signal is given (at least 5% activation), which also narrows down the sampling to parameter values for which the temporal ultrasensitivity can reliably be determined. The resulting selection of parameter samples is denoted as ‘Pre’ in this figure. The subsequent step is to select for parameter samples that result in temporal ultrasensitivity higher than 0.2, yielding the ‘Post’ parameter collection. Comparison of the parameter values found in the Pre and Post collections gives information about the preference for certain parameter values for circuits with high temporal ultrasensititivity. B) The distribution of the values of each circuit parameter in the Pre and Post collections of parameter sets visualized as box plots with a dashed green line indicating the parameter value obtained from a fit to the experimental data. For most parameters, the Pre and Post distributions are comparable, meaning that those parameter values are equally likely to be found before and after selection for a high temporal ultrasensitivity. The parameters that display the largest shift in distribution between Pre and Post are $bgratio_{sw}$, the fraction the translation rate of the bound switch:trigger complex that is observed as leakage in the unbound toehold switch, and $N_{S28}$, which is the Hill coefficient of the binding of $\sigma^{28}$ to its promoter. A high temporal ultrasensitivity is associated with a lower $bgratio_{sw}$ and a higher $N_{S28}$. C) 2D distributions of parameter values of a select combination of parameters, displaying the Pre and Post collections. These plots show which combinations of parameter values are enriched when selecting for a high temporal ultrasensitivity.

| Figure | Type | Samples | DNA | DNA Name | Concentration (nM) |
|---|---|---|---|---|---|
| 1c | Batch | All | | P70a-SwitchA-eGFP | 2 |
| | | On-Target | X1/X3 | P70a-TriggerA | 10 |
| | | Off-Target | | P70a-TriggerC | 10 |
| 1d | Batch | All | Y1/Y3 | P70a-SwitchA-S28 | 1 |
| | | All | Z1 | P28a-SwitchA-eGFP | 10 |
| | | On-Target | X1/X3 | P70a-TriggerA | 10 |
| | | Off-Target | X2 | P70a-TriggerB | 10 |
| 2b | Batch | CFFL 1: All | Y1/Y3 | P70a-SwitchA-S28 | 1 |
| | | CFFL 2: All | Y2 | P70a-SwitchB-S28 | 0.6 |
| | | CFFL 3: All | Y1/Y3 | P70a-SwitchA-S28 | 0.8 |
| | | CFFL 1: All | Z1 | P28a-SwitchA-eGFP | 10 |
| | | CFFL 2: All | Z2 | P28a-SwitchB-eGFP | 10 |
| | | CFFL 3: All | Z3 | P28a-SwitchA-eCFP | 10 |
| | | CFFL 1 / 3: On-Target | X1/X3 | P70a-TriggerA | 10 |
| | | CFFL 1: Off-Target | | P70a-TriggerC | 10 |
| | | CFFL 3: Off-Target | X2 | P70a-TriggerB | 10 |
| | | CFFL 2: On-Target | X2 | P70a-TriggerB | 10 |
| | | CFFL 2: Off-Target | X1/X3 | P70a-TriggerA | 10 |
| 2d | Batch | All | Y2 | P70a-SwitchB-S28 | 0.6 |
| | | All | Y1/Y3 | P70a-SwitchA-S28 | 0.8 |
| | | All | Z2 | P28a-SwitchB-eGFP | 10 |
| | | All | Z3 | P28a-SwitchA-eCFP | 10 |
| | | Full / -X2 | X1/X3 | P70a-TriggerA | 10 |
| | | Full / -X3 | X2 | P70a-TriggerB | 10 |
| 2e | Batch | All | Z2 | P28a-SwitchB-eGFP | 10 |
| | | All | Z3 | P28a-SwitchA-eCFP | 10 |
| | | Full / -Y2 / -X2 -Y2 | Y1/Y3 | P70a-SwitchA-S28 | 0.8 |
| | | Full / -Y3 / -X3 - Y3 | Y2 | P70a-SwitchB-S28 | 0.6 |
| | | Full / -Y2 | X1/X3 | P70a-TriggerA | 10 |
| | | Full / -Y3 | X2 | P70a-TriggerB | 10 |
| 3b | Batch | All | Y1/Y3 | P70a-SwitchA-S28 | 0.1, 0.3, 0.5, 1, 2, 5 |
| | | All | Z1 | P28a-SwitchA-eGFP | 10 |
| | | *ON* State | X1/X3 | P70a-TriggerA | 10 |
| 3c | Batch | All | Y1/Y3 | P70a-SwitchA-S28 | 5 |

| | | | | | |
|---|---|---|---|---|---|
| | | All | Z1 | P28a-SwitchA-eGFP | 1, 2, 5, 10, 20 |
| | | *ON* State | X1/X3 | P70a-TriggerA | 10 |
| 3d | Batch | CFFL 2 (solid green): All / Composite: All | Y2 | P70a-SwitchB-S28 | 0.6 |
| | | CFFL 3 (solid blue): All / Composite: All | Y1/Y3 | P70a-SwitchA-S28 | 0.8 |
| | | CFFL 2 (solid green): All / Composite: All | Z2 | P28a-SwitchB-eGFP | 10 |
| | | CFFL 3 (solid blue): All / Composite: All | Z3 | P28a-SwitchA-eCFP | 10 |
| | | CFFL 2 (solid green): *ON* State / Composite left (dashed green): *ON* State | X2 | P70a-TriggerB | 10 |
| | | CFFL 3 (solid blue): *ON* State / Composite right (dashed blue): *ON* State | X1/X3 | P70a-TriggerA | 10 |
| 4b | Flow | All: *ON* State | X1/X3 | P70a-TriggerA | 10 |
| | | All: All | Z1 | P28a-SwitchA-eGFP | 10 |
| | | CFFL: All | Y1/Y3 | P70a-SwitchB-S28 | 0.2, 0.5, 1, 5 |
| | | Reference Motif: All | R1 | P70a-S28 | 0.3 |
| 4c | Flow | *ON* State | X1/X3 | P70a-TriggerA | 10 |
| | | All | Z1 | P28a-SwitchA-eGFP | 10 |
| | | All | Y1/Y3 | P70a-SwitchB-S28 | 1 |
| | | Input durations: 15, 30, 60, 120, persistent | | | |
| 5a | Flow | All: *ON* State | X1/X3 | P70a-TriggerA | 10 |
| | | All: All | Z1 | P28a-SwitchA-eGFP | 10 |
| | | CFFL: All | Y1/Y3 | P70a-SwitchB-S28 | 0 |
| | | Reference Motif: All | R1 | P70a-S28 | 0.3 |
| | | Input durations: 0, 15, 30, 60, 120, persistent | | | |
| 5d/e | Model | All: *ON* State | X1/X3 | P70a-TriggerA | 10 |
| | | All: All | Z1 | P28a-SwitchA-eGFP | 10 |
| | | CFFL: All | Y1/Y3 | P70a-SwitchB-S28 | $10^{-3}$ - $10^{2}$ |
| | | Reference Motif: All | R1 | P70a-S28 | $10^{-3}$ - $10^{2}$ |
| | | Input durations: logspace(-2,1.3,100) | | | |
| 6d | Model | All | X1/X3 | P70a-TriggerA | 10 |

| | | | | | |
|---|---|---|---|---|---|
| | | All: All | Z1 | P28a-SwitchA-eGFP | 10 |
| | | CFFL: All | Y1/Y3 | P70a-SwitchB-S28 | 1 |
| | | Reference Motif: All | R1 | P70a-S28 | 1 |
| | | Input durations: $10^{-2}$ - $10^{2}$ | | | |
| S2 | Batch | Initial: All | | P70a-SCAR-SwitchA-eGFP | 10 |
| | | Optimized: All | | P70a-SwitchA-eGFP | 10 |
| | | All: +Trigger | X1/X3 | P70a-TriggerA | 20 |
| S3a | Batch | All | | P70a-eGFP | 0.01, 0.02, 0.05, 0.1, 0.2, 0.5, 1, 2, 5, 10, 20 |
| S3b | Batch | Left (orange) | | P70a-SwitchA-eGFP | 2 |
| | | Left (orange) | X1/X3 | P70a-TriggerA | 1, 2, 5, 6, 10, 20 |
| | | Right (green) | X1/X3 | P70a-TriggerA | 10 |
| | | Right (green) | | P70a-SwitchA-eGFP | 1, 2, 5, 10 |
| S4 | Batch | All | | P70a-eGFP | 2 |
| | | All | R1 | P70b-S28 | 0.5, 1, 2, 5, 10 |
| S5a | Batch | On-target | X1/X3 | P70a-TriggerA | 10 |
| | | Off-target | | P70a-TriggerC | 10 |
| | | All | | P28a-eGFP | 5 |
| | | All | Y1/Y3 | P70a-SwitchB-S28 | 0.1 |
| S5b | Batch | All | X1/X3 | P70a-TriggerA | 10 |
| | | All | | P28a-eGFP | 10 |
| | | All | Y1/Y3 | P70a-SwitchB-S28 | 0.1, 0.2, 0.5, 1, 2, 5, 10 |
| S6b | Batch | All | R1 | P70a-S28 | 5 |
| | | All | Z1 | P28a-SwitchA-eGFP | 10 |
| | | On-Target | X1/X3 | P70a-TriggerA | 10 |
| | | Off-Target | X2 | P70a-TriggerB | 10 |
| S7 | Batch | All | X1/X3 | P70a-TriggerA | 10 |
| | | All | X2 | P70a-TriggerB | 10 |
| | | Full / -Y3 -Z2 / -Y2 -Y3 | Z3 | P28a-SwitchA-eCFP | 10 |
| | | Full / -Y2 -Z3 / -Y2 -Y3 | Z2 | P28a-SwitchB-eGFP | 10 |
| | | Full / -Y3 -Z2 | Y2 | P70a-SwitchB-S28 | 0.6 |
| | | Full / -Y2 -Z3 | Y1/Y3 | P70a-SwitchA-S28 | 0.8 |

| | | | | | |
|---|---|---|---|---|---|
| S8 | Batch | P70a/b: All | Z1 | P28a-SwitchA-eGFP | 10 |
| | | P70a: All | R1 | P70a-S28 | 0.1, 0.5, 1, 2, 5 |
| | | P70b: All | R1 | P70b-S28 | 0.5, 1, 2, 5, 5, 10 |
| | | P70a/b: *ON* State | X1/X3 | P70a-TriggerA | 10 |
| S9a/b | Model | All: *ON* State | X1/X3 | P70a-TriggerA | 10 |
| | | CFFL: All | Y1/Y3 | P70a-SwitchB-S28 | 1 |
| | | Reference Motif: All | R1 | P70a-S28 | 1 |
| | | All: All | Z1 | P28a-SwitchA-eGFP | 10 |

**Table S3:** DNA construct concentrations of all batch, flow and *in silico* experiments.

| Experiment | Sample | DNA ***X1*** (nM) | DNA ***Y1*** (nM) | DNA ***R1*** (nM) | DNA ***Z1*** (nM) |
|---|---|---|---|---|---|
| CFFL, ***Y1*** = 0.2 nM | *No input* | 0 | 0.57 | 0 | 29 |
| | *High input* | 71 | 0.57 | 0 | 29 |
| | *Low input* | 29 | 0.57 | 0 | 29 |
| CFFL, ***Y1*** = 0.5 nM | *No input* | 0 | 1.4 | 0 | 29 |
| | *High input* | 71 | 1.4 | 0 | 29 |
| | *Low input* | 29 | 1.4 | 0 | 29 |
| CFFL, ***Y1*** = 1 nM | *No input* | 0 | 2.9 | 0 | 29 |
| | *High input* | 71 | 2.9 | 0 | 29 |
| | *Low input* | 29 | 2.9 | 0 | 29 |
| CFFL, ***Y1*** = 5 nM | *No input* | 0 | 14 | 0 | 29 |
| | *High input* | 71 | 14 | 0 | 29 |
| | *Low input* | 29 | 14 | 0 | 29 |
| Reference Motif, ***R1*** = 0.3 nM | *No input* | 0 | 0 | 0.86 | 29 |
| | *High input* | 71 | 0 | 0.86 | 29 |
| | *Low input* | 29 | 0 | 0.86 | 29 |

**Table S4:** Composition of the DNA solutions used in flow experiments. A flow experiment of an initial fill of the reactor with TXTL reaction mixture and 35% reactor volume *No input* solution. Subsequently, every 15 minutes 40% of the reactor was refreshed with a mixture consisting of 65% TXTL mixture and the remaining 45% one of the DNA solutions. The sequence of DNA solutions was: 11 or 15 steps *No input*; 1 step *High input*; 0, 1, 3 or 7 steps *Low input* (to create 15 min, 30 min, 1 h and 2 h input pulses) and the remaining steps *No input* until 11 h of reaction time were reached. Additionally, a negative control without input signal was constructed by only supplying the *No input* solution. The persistent input experiments were conducted using the following sequence: 11 or 15 steps *No input*; 1 step *High input* and the remaining steps *Low input*.

### References

(1) Sun, Z. Z., Yeung, E., Hayes, C. A., Noireaux, V., and Murray, R. M. (2014) Linear DNA for Rapid Prototyping of Synthetic Biological Circuits in an *Escherichia coli* Based TX-TL Cell-Free System. *ACS Synthetic Biology 3*, 387–397.

(2) Yelleswarapu, M., van der Linden, A. J., van Sluijs, B., Pieters, P. A., Dubuc, E., de Greef, T. F. A., and Huck, W. T. S. (2018) Sigma Factor-Mediated Tuning of Bacterial Cell-Free Synthetic Genetic Oscillators. *ACS Synth. Biol. 7*, 2879–2887.

(3) Gerardin, J., Reddy, N. R., and Lim, W. A. (2019) The Design Principles of Biochemical Timers: Circuits that Discriminate between Transient and Sustained Stimulation. *Cell Systems 9*, 297-308.e2.
